# The brain’s topographical organization shapes dynamic interaction patterns to support flexible behavior

**DOI:** 10.1101/2023.09.06.556465

**Authors:** Xiuyi Wang, Katya Krieger-Redwood, Baihan Lyu, Rebecca Lowndes, Guowei Wu, Nicholas E. Souter, Xiaokang Wang, Ru Kong, Golia Shafiei, Boris C. Bernhardt, Zaixu Cui, Jonathan Smallwood, Yi Du, Elizabeth Jefferies

**Affiliations:** CAS Key Laboratory of Behavioral Science, Institute of Psychology, Chinese Academy of Sciences, Beijing, 100101, China; Department of Psychology, University of Chinese Academy of Sciences, Beijing 100049, China; Department of Psychology, University of York, Heslington, York, YO10 5DD, United Kingdom; Department of Biomedical Engineering, University of California, Davis, CA 95616, USA; Department of Electrical and Computer Engineering, National University of Singapore, Singapore, Singapore; Department of Psychiatry, Perelman School of Medicine, University of Pennsylvania, Philadelphia, PA 19104, USA; McConnell Brain Imaging Centre, Montreal Neurological Institute and Hospital, McGill University, Montreal, QC, Canada; Chinese Institute for Brain Research, Beijing 102206, China; Department of Psychology, Queens University, Kingston, Ontario, Canada; CAS Center for Excellence in Brain Science and Intelligence Technology, Shanghai 200031, China

**Author notes:** Author contributions: X.W., Y.D. and E.J. designed research; K.K.R., R. L. and X.W. collected the data, X.W., B.L., and G.W. analyzed data; K.K.R., Z.C., X.W., provided suggestions about data analysis; X.W., Y.D. and E.J. wrote the original manuscript. All authors edited the manuscript.

**Keywords:** flexible cognition, working memory, long-term memory, fronto-parietal control network, dorsal attention network, default mode network, cortical topography, cortical hierarchy

## Abstract

Understanding how human cognition flexibly supports distinct forms of behavior is a key goal of neuroscience. Adaptive behavior relies on context-specific rules that vary across situations, as well as on stable knowledge gained from experience. However, the mechanisms that allow these influences to be appropriately balanced remain elusive. Here, we show that this cognitive flexibility is partly supported by the topographical organization of the cortex. The frontoparietal control network (FPCN) is located between regions implicated in top-down attention and memory-guided cognition. We hypothesized that the FPCN is topographically divided into discrete systems that support these distinct forms of behavior. These FPCN subsystems exhibit multiple anatomical and functional similarities to their neighboring systems (the dorsal attention network and default mode network respectively). This topographic architecture is also mirrored in the functional patterns that emerge in different situations: the FPCN subnetworks act as a unified system when long-term memory can support behavior, but they segregate into discrete units when working memory, rather than long term memory, is necessary for behavioral control. In this way, the topographic organization of brain function provides crucial insights into how the human cortex supports flexible behavior.

## Introduction

Human behavior is inherently flexible, with effective actions varying substantially across different situations. In some circumstances, behavior must be guided by information in working memory, which allows for the temporary and effortful maintenance of rules that are relevant to the current situation. In other scenarios, behaviors are guided by information from long-term memory, such as our general knowledge of the world gained through a lifetime of experience ^1,2^. These two modes of cognition depend on different neural resources and at times they can be antagonistic. For example, situations like mind-wandering illustrate what can happen when the contextual regulation of cognition fails ^3,4^. As a result, a key question in neuroscience is how the human brain achieves an appropriate functional balance between these two fundamental cognitive modes in a situation specific manner to support flexible behavior.

Research has established that behaviors guided by external inputs encoded in working memory often rely on neural processes within the dorsal attention network (DAN) ^5^, while situations in which long-term memory supports cognition often draw on the default mode network (DMN) ^6^. It is hypothesized that these influences are balanced by the brain’s control system — the fronto-parietal control network (FPCN) ^7–11^. Regions of the FPCN are widely distributed across the frontal and parietal lobes (see Fig. 1A) and are proposed to implement cognitive control by dynamically coordinating activity among diverse brain systems to integrate brain-wide processing in a goal directed manner ^7–11^. Despite evidence that the FPCN is commonly recruited across tasks ^12–16^, it remains unclear how a single system can flexibly support the distinct modes of operation required for the wide range of human behaviors. Contemporary work suggests that the ability to flexibly adjudicate between working memory and long-term knowledge as drivers of behavior might be achieved by subdividing the control network into two topographically adjacent yet distinct subnetworks FPCN-A and FPCN-B (see Fig. 1A). It is hypothesized that regions of FPCN-A, which are closer to sensory-motor regions, are linked to behavioral control when optimal behavior depends on working memory information, such as task rules, while FPCN-B, which is more anterior on the cortical mantle, is linked to situations when memory is essential for adaptive behavioral control (see Fig. 1A). Consistent with this perspective, FPCN subnetworks are hypothesized to have different patterns of activation and functional connectivity: FPCN-A exhibits stronger functional connectivity with DAN, while FPCN-B exhibits stronger functional connectivity with DMN ^11,17,18^. Our study builds on this emerging evidence to ask (i) how underlying anatomical differences within the FPCN relate to its functional differentiation into subsystems, and (ii) how these topographically separated systems adapt their interaction patterns to interface with anti-correlated networks (DAN and DMN) in a way that supports multiple different modes of operation that allow flexible behavior ^11,17,18^.

Our study tests the emerging hypotheses that subnetworks of FPCN are topographically proximal to systems linked to external attention and to long-term knowledge, which are differentiated anatomically, and that this differentiation enables them to interact with spatially adjacent systems. This proposal is related to emerging evidence that the geometry of the cortex (i.e., the shape) sculpts its functions ^19^ and the observation that the principal dimension describing functional differentiation within the cortex, often referred to as the principal gradient, outlines the physical sequence of networks on the cortical surface ^20^. This principal dimension of functional connectivity within the cortex is anchored at one end by the default mode network and at the other by sensory systems. In this topographical scheme, the FPCN is located between these two extremes. Compared with sensory-motor systems, which are located at one end of a cortical hierarchy, DMN regions at the other end show lower myelination ^22,24,25^, the lowest correspondence between structure and function ^26,27^, the lowest functional similarity across species, and the greatest cortical expansion from macaque to human ^28,29^. Given these observations, we hypothesized that FPCN-A and FPCN-B share more similar anatomical features with DAN and DMN, respectively. Yet although the broad topographical patterns spanning from unimodal to transmodal regions are well documented ^1,6,20,21^, the patterns for different measurements are not identical ^20,22,23^, leaving the precise topographical positions of FPCN subsystems an open question. Furthermore, a novel contribution of this study is to contextualize the mechanisms that underpin FPCN-A and FPCN-B’s distinctive functional roles within this anatomical framework, helping to explain neural and cognitive flexibility.

We investigate how the topography of FPCN subsystems relates to their functional interactions with other networks. The FPCN subsystems were defined using Kong’s parcellation (2021) ^30^, which subdivides the FPCN into subnetworks that capture the inherent functional variability of the FPCN. This method enabled us to generate individual-specific parcellations with heightened homogeneity, allowing us to depict brain organization more accurately. We employed well-designed tasks to establish the parcels’ functional differentiation and to uncover how their interaction patterns change across contexts to support flexible behavior. We hypothesize that the relative proximity of FPCN-A to DAN, and of FPCN-B to DMN, plays a pivotal role in flexibly generating distinct cognitive modes that are relevant to the updating of working memory versus retrieval from long-term memory (Fig. 1). Critically, while this topographical organization suggests that adjacent networks will have more similarity, both structurally and functionally, the brain also produces flexible patterns of behavior based on task demands, in which different subsets of adjacent networks are recruited together to address either external or internal task requirements. By examining functional similarity across tasks states, we can establish how topography supports different cognitive modes, while also confirming that patterns of similarity are not solely a consequence of network adjacency. In this way, we establish how the topographical organization of the cortical mantle enables diverse interactions of FPCN with networks linked to top-down attention and long-term memory, giving rise to different landscapes of neural activity in response to different situational demands.

**Fig. 1.**
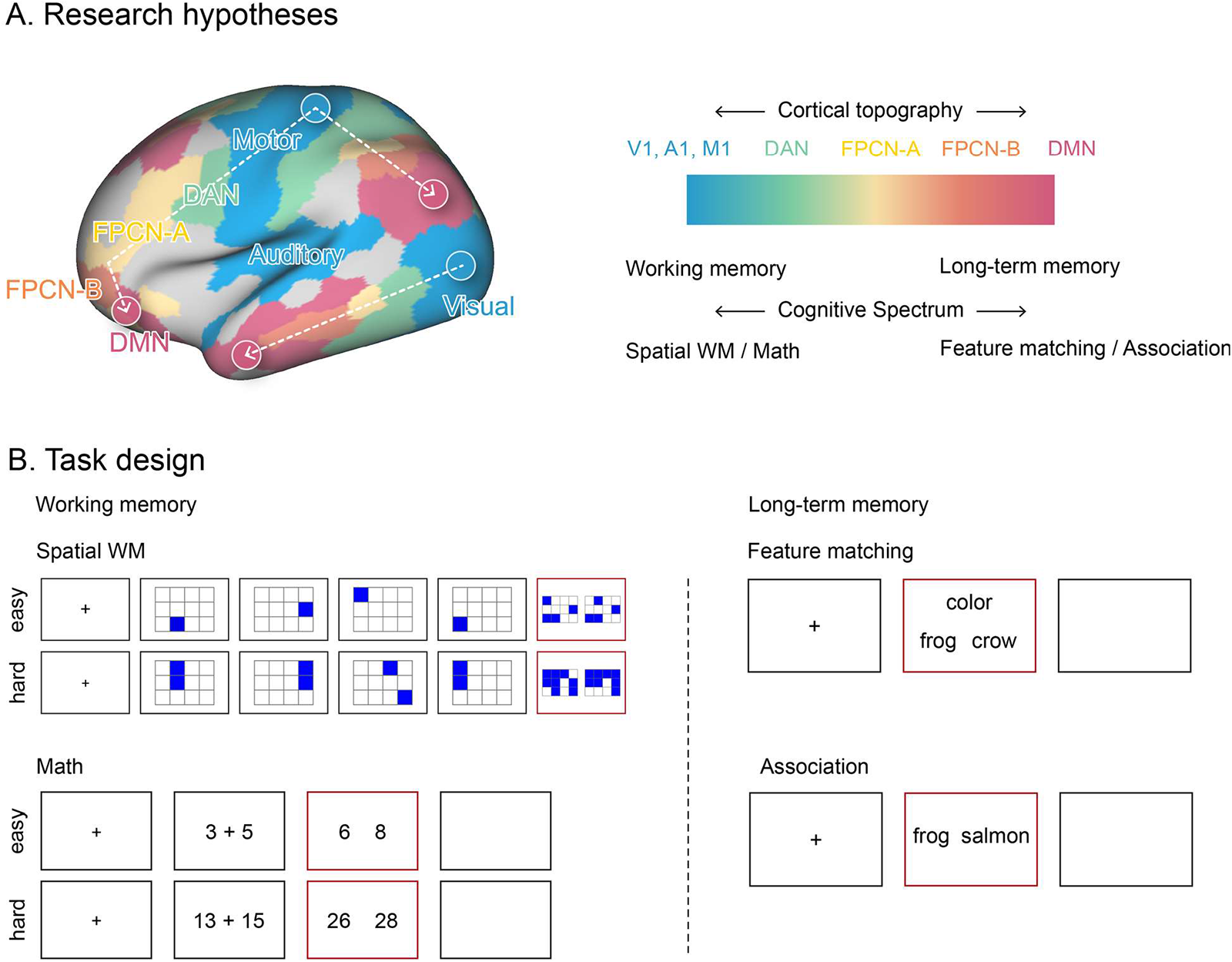
The research hypothesis and task design. A: The research framework. Brain organization relates to cortical topography: brain regions supporting perceptual/motor features are maximally separated from heteromodal aspects of long-term memory, with control regions in the middle. This key dimension of topographical organization might relate to a ‘cognitive spectrum’ capturing distinctions between tasks that rely on recently presented information in working memory versus long-term knowledge (other spectrums may also exist). Regions of FPCN-A are closer to sensory-motor regions and more posterior than those of FPCN-B on the cortical mantle. These networks were identified using Kong’s parcellation approach ^30^, which generates individual-specific parcellations with greater homogeneity. Other networks, such as DAN and DMN, are also divided into subnetworks. The subnetworks of visual, motor, DAN, and DMN are merged here for illustration. See Fig 3 for the distribution of each subnetwork. B: Task design. To tap working memory, we included two tasks: a spatial working memory task required participants to keep track of sequentially presented locations, while math decisions involved maintaining and manipulating numbers which rely more on working memory. To tap long-term memory, we included two tasks that required controlled retrieval of knowledge; a semantic feature matching task required participants to match probe and target concepts according to a particular semantic feature (color or shape), while a semantic association task involved deciding if pairs of words were linked in meaning. Response periods are indicated by a red box. (V1 = Visual, A1 = Auditory, M1 = Motor, DAN = Dorsal attention network, FPCN = Fronto-parietal control network, DMN = Default mode network).

## 2. Results

The results are divided into three sections: (i) first, we take an existing individualized parcellation of the cortex identifying two large-scale distributed control networks, corresponding to FPCN-A and FPCN-B, and establish that parcels of these networks have reliably different topographical locations by quantifying their distance from sensory-motor regions on multiple metrics; (ii) next we ask how these differences in topography of FPCN-A and FPCN-B relate to their functional interaction patterns; (iii) finally, we demonstrate how these interaction patterns produce flexible behavior across different task contexts.

We selected Kong et al.’s parcellation (2021) ^30^ as the most appropriate choice, since this parcellation subdivides the FPCN into subnetworks, which is essential given the inherent heterogeneity of the FPCN, and because this method enables us to generate individual-specific parcellations with heightened homogeneity. The naming of FPCN-A and FPCN-B here was consistent with the original Yeo et al.’s parcellation (2011) ^31^ and Kong et al.’s individualized parcellation ^30^, adapted from the Yeo et al.’s parcellation. However, the opposite naming has been used in some previous studies that investigate the functional differentiation of FPCN ^17,18^. We did not use other parcellations, including Power et al. (2011) ^32^, Gordon et al. (2016) ^33^, and Glasser et al. (2016) ^24^ etc., because FPCN was identified as a functional unit without further subdivisions in these atlases.

### 2.1 The topographical characteristics of FPCN-A and FPCN-B

To establish whether the FPCN-A subnetwork is closer to sensory-motor systems, while FPCN-B is proximal to DMN, we examined the topographical positions of FPCN-A and FPCN-B on multiple metrics, including cortical geometry, anatomical characteristics, principal connectivity gradient values, cortical expansion, and cross-species functional similarity. We examined multiple metrics because although these characteristics generally show similar topographical patterns – spanning from unimodal to transmodal ^1,6,20,21^ – they do not perfectly align with each other. For example, the correspondence between structure and function is weak in the transmodal area ^26^, and the minimum physical distance to the sensory-motor landmarks shows only a moderate correlation with the principal connectivity gradient ^20^. These observations suggest a degree of dissociation between these metrics, leaving the precise topographical positions of FPCN subsystems as an open question. This motivated us comprehensively capture the topographic positioning of FPCN subnetworks across multiple metrics.

#### 2.1.1 FPCN-A is physically closer to sensorimotor cortex than FPCN-B

Distance from sensorimotor regions is thought to provide an organizing principle of functional differentiation within the cortex ^1,6,21,34^. Therefore, our first analysis confirmed that there were systematic differences in the location of the FPCN-A and FPCN-B networks on the cortical surface, in terms of their physical distance to primary sensory-motor landmarks, using the human connectome project (HCP) dataset. We calculated the geodesic distance between each parcel and three key landmarks associated with primary visual, auditory and somatomotor cortices to identify the global minimum geodesic distance to primary sensorimotor regions for each parcel. Fig. 2A shows a group-level representation of global minimum distance from sensory-motor cortex: transmodal regions are further from these landmarks. FPCN-A showed greater physical proximity to sensorimotor cortex than FPCN-B (t = -100.57, p < 0.001; Fig. 2A and 2B).

**Fig. 2.**
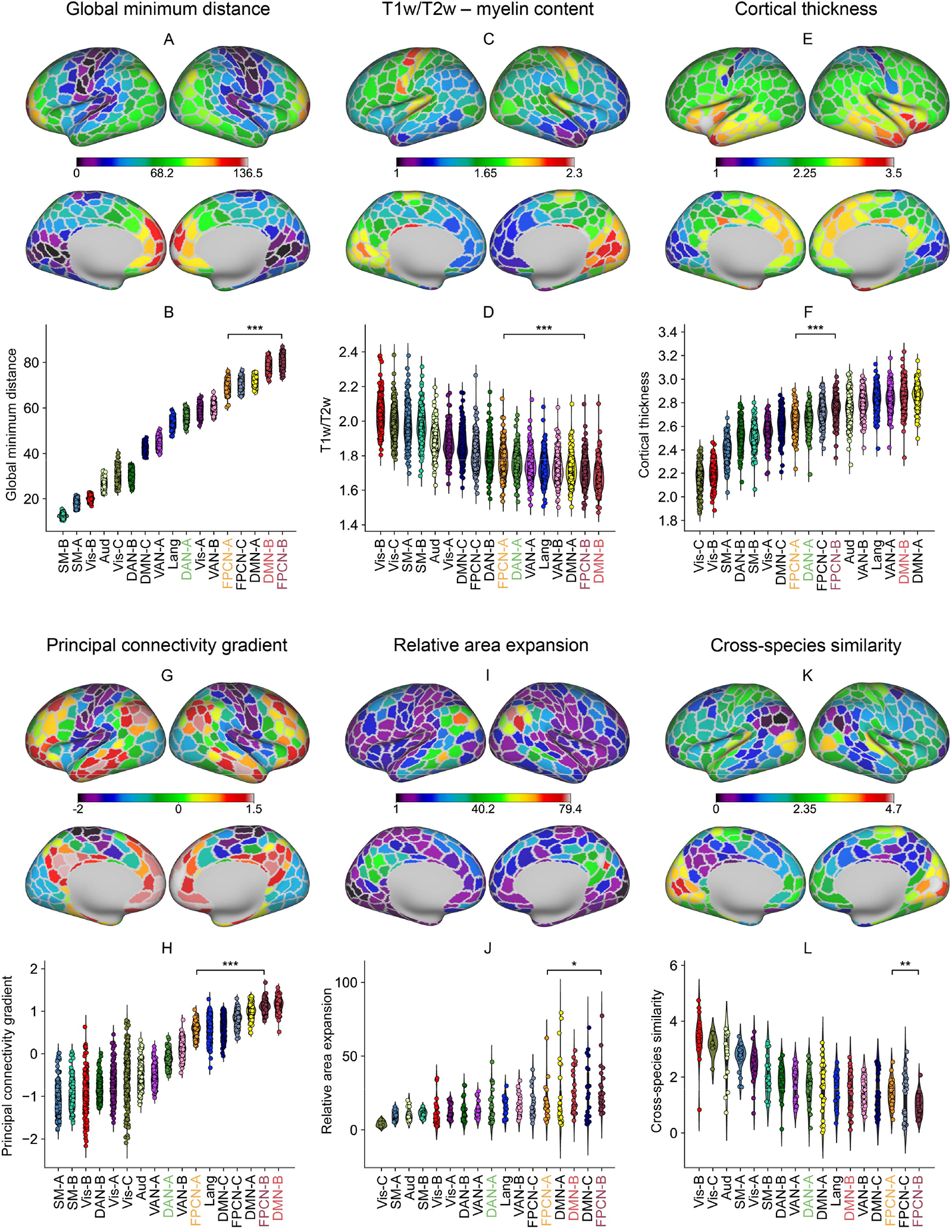
FPCN-A was closer to DAN and sensory-motor systems, while FPCN-B was proximal to DMN in physical distance, myelin content, cortical thickness, principal connectivity gradient values, cortical expansion, and cross-species in functional similarity. Each datapoint in B, D, F, and H represents the data of one participant and each datapoint in J and L represents the data of one parcel. We sorted the networks by their mean values across participants in B, D, F, and H and across parcels in J and L. A and B – The global minimum geodesic distance between each parcel and its closet sensory-motor landmark and the global minimum distance for each network (averaged across parcels). FPCN-A was closer to sensory-motor landmarks than FPCN-B. C and D – T1w/T2w values were significantly lower in association networks than in sensory networks. The T1w/T2w value in FPCN-A was higher than in FPCN-B and was closer to the value for sensory-motor cortex. E and F – Cortex was generally thicker in control networks and DMN than in sensory motor networks. The cortex in FPCN-A was thinner than in FPCN-B. G and H – The principal connectivity gradient that explained the most variance in resting-state fMRI captured the separation between sensory-motor and transmodal regions. FPCN-A was closer to the sensory-motor end of this gradient axis than FPCN-B. I and J – Sensory cortices expanded the least from the macaque to the human, while transmodal cortex expanded the most. FPCN-A showed less cortical expansion than FPCN-B. K and L – Regions of sensory-motor networks showed greater cross-species similarity between humans and macaques, whereas transmodal regions showed greater differences. FPCN-A showed greater cross-species similarity than FPCN-B.

#### 2.1.2 FPCN-A was closer than FPCN-B to the unimodal end of the anatomical organization defined by myelin content and cortical thickness

Having confirmed topographic differences in the control subnetworks, we next examined whether these are mirrored in anatomical differences. The cortical T1w/T2w map – sensitive to regional variation in grey-matter myelin content ^22,24,35^ – is thought to reflect an anatomical hierarchy, with sensorimotor regions showing greater myelination ^22^ . We hypothesised that FPCN-A would have higher levels of myelination than FPCN-B since the analysis above showed that FPCN-A was closer to sensorimotor cortex. We analysed participants’ individual T1w/T2w maps and cortical thickness maps from the HCP dataset. As expected, myelin values were high in sensorimotor cortices and low in association cortices. Among the transmodal networks, FPCN-A had higher myelin values than FPCN-B (t = 46.89, p < 0.001; Fig. 2C and 2D). In parallel, since cortical thickness coarsely tracks changes in cytoarchitecture and myelin content, we found that the cortex was generally thicker in heteromodal control networks and DMN than in sensory motor networks (Fig. 2E and 2F). FPCN-A had lower cortical thickness than FPCN-B (t = 34.08, p < 0.001).

#### 2.1.3 FPCN-A was closer to the unimodal end of the principal connectivity gradient than FPCN-B

Having established that anatomical differences reflect topographical proximity, we next explored whether resting state functional connectivity shows a similar pattern. Global minimum distance is positively correlated with location on the principal connectivity gradient, which organizes neural systems along a spectrum from unimodal to transmodal cortex ^20,36^. We therefore asked whether the parcels of FPCN-A were closer than FPCN-B to the unimodal end of the principal connectivity gradient. Dimension reduction analysis was performed on the HCP resting state functional connectivity matrix. For 238 out of 245 participants, the dimension explaining the most variance corresponded to the principal gradient as described by Margulies et al. (2016) ^20^: sensory-motor regions fell at one end of this dimension of connectivity (shown in purple-blue in Fig. 2G), while transmodal areas were located at the other end (shown in red-orange in Fig. 2H). We averaged the principal gradient values of all the parcels within each network for all participants for whom the principal gradient explained the most variance. We found that sensory-motor networks fell at one end, while control networks and DMN were located the other end. FPCN-A had lower values on the principal gradient than FPCN-B (t = 51.93, p < 0.001; Fig. 2G and 2H), indicating that FPCN-A was closer to sensorimotor systems on this dimension of connectivity, while FPCN-B was closer to the DMN apex of the principal gradient.

#### 2.1.4 FPCN-A showed less cortical expansion from macaque to human and showed greater similarity across species relative to FPCN-B

A prominent theory of cortical organization suggests that transmodal networks became untethered from sensorimotor systems through evolution ^21^. Therefore, we hypothesized that FPCN-B would show more cortical expansion and less cross-species similarity in functional connectivity than FPCN-A, since it was further from sensorimotor systems. We used the evolutionary expansion and cross-species similarity map provided by Xu et al .^29^. To estimate surface areal expansion, human surface area was divided by macaque surface area at each of corresponding vertex and then all vertices within each parcel were averaged. We found that FPCN-B showed more expansion (t = 2.125, p = 0.039; Fig. 2I and 2J) and less cross-species functional similarity than FPCN-A (t = -3.333, p = 0.002; Fig. 2K and 2L).

**Fig. 3.**
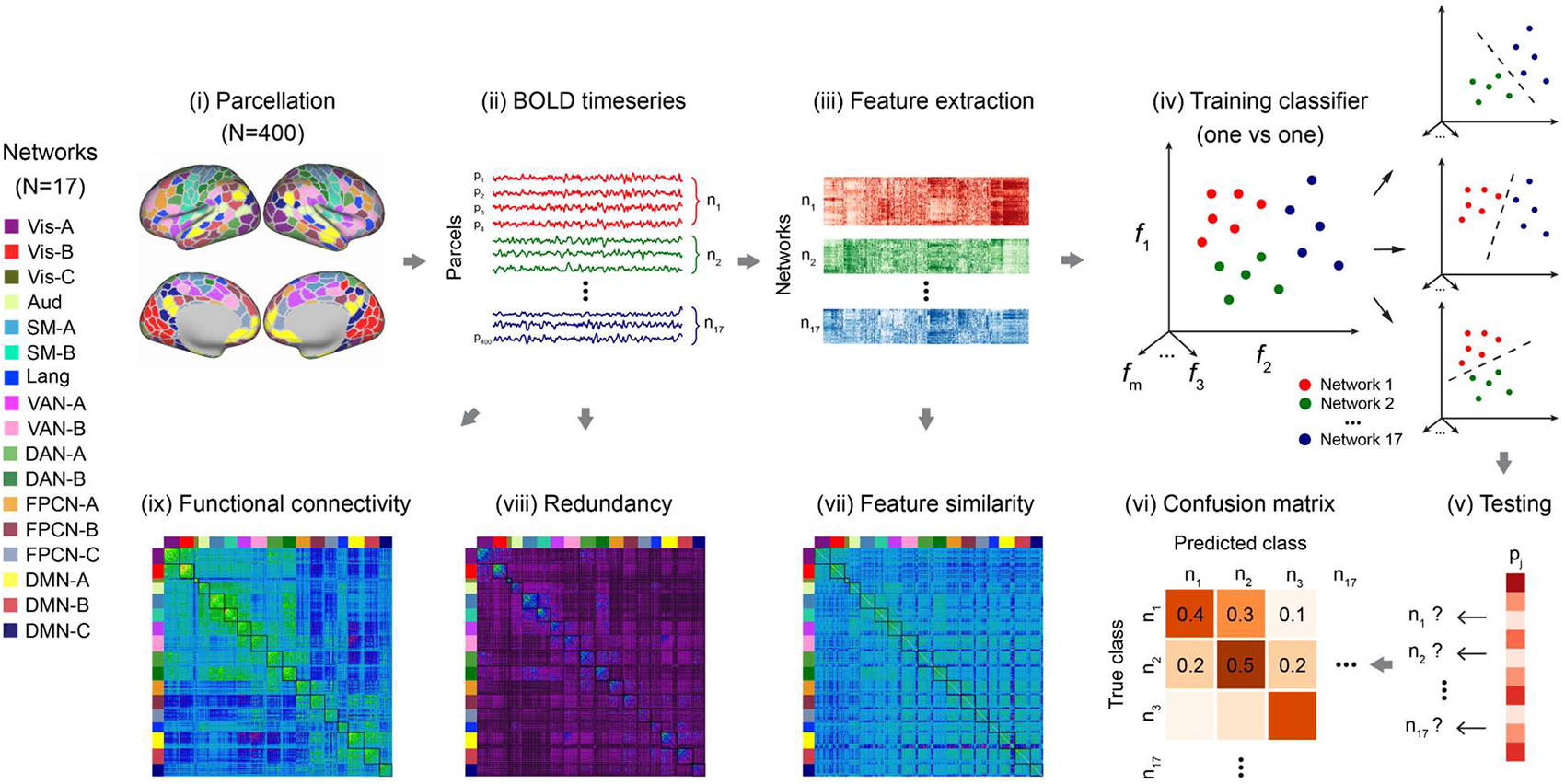
Overview of analytic approaches to study the functional interaction patterns of brain networks. (I) Individual-specific parcellation divided the whole brain into 400 parcels across 17 networks ^30^. (II) Average time series of each parcel. (III) Extraction of features of time series for each parcel. (IV) Multi-class classification analysis involved training a classifier to learn the mapping from time-series features of parcels to network labels and (V) then to predict network labels for parcels. (VI) Network similarity was characterized by the normalized confusion matrix: networks with more similar functions would be more likely to be incorrectly classified as each other. (VII) Pearson correlation coefficients of the extracted features represent the pairwise feature similarity between all possible combinations of brain parcels. (VIII) Redundancy quantifies how much information about the brain’s future trajectory is predicted redundantly by distinct brain regions. We focused on temporally persistent redundancy, which corresponds to redundant information in the past of both regions that is also present in the future ^37^. (IX) Functional connectivity involved calculating Pearson correlation coefficients between the time-series of parcels. Vis = Visual, Aud = Auditory, SM = Sensory-motor, DAN = Dorsal attention network, VAN = Ventral attention network, FPCN = Fronto-parietal control network, Lang = Language, DMN = Default mode network, WM = working memory.

### 2.2 Functional interaction and activation patterns of FPCN-A and FPCN-B

The results above show that FPCN-A and FPCN-B occupy distinct topographical positions: FPCN-A is closer to sensory-motor landmarks and DAN, while FPCN-B is closer to DMN in geodesic distance, anatomical features relating to myelination, functional connectivity patterns, and evolutionary markers. Since DAN and DMN typically show negative functional connectivity ^38^, control subnetworks that are proximal to these systems may show a degree of functional separation that reflects their topography. To investigate this possibility, we investigated multiple metrics of functional similarity, comparing DAN and DMN with control subnetworks A and B to establish whether they showed dissociable patterns of functional recruitment in rest, working memory and long-term memory tasks. We would expect greater functional similarity between FPCN-A and adjacent DAN, and between FPCN-B and adjacent DMN if topography constrains brain function^1,6,20,39,40^. We investigated this prediction in three complementary analyses examining (i) feature classification across networks, (ii) functional coupling, and (iii) activation and deactivation patterns modulated by task difficulty. Since no single method can conclusively pinpoint network interactions ^37,41–43^, comprehensive evidence from various methods, when contextualized within our broader understanding of brain anatomy and function, will provide the most reliable insights.

#### 2.2.1 FPCN-A was more likely to be misclassified as DAN-A and FPCN-B was more likely to be misclassified as DMN-B

To reveal network similarity, multi-class classification was used to predict the network labels of parcels using extracted features of the time-series data. These features include temporal autocorrelation, kurtosis, and entropy ^44^ etc., which may capture meaningful differences between different types of time series and thus represent promising candidates as quantitative phenotypes for distinguishing data of different types (see Method 4.6.6 for feature extraction, Method 4.6.7 for classification analysis, Fig. 3 for the analysis pipeline). This analysis was motivated by the observation that certain brain regions possess distinct features, which equip them to support different functions, while regions with similar features are suited to support similar functions, irrespective of whether they exhibit positive or negative functional connectivity. For instance, early visual areas and the DMN are situated at opposing ends of the timescale hierarchy. The former boasts the shortest timescale, marked by rapid temporal autocorrelation decay, while the latter has the longest, characterized by gradual autocorrelation decay ^23,45–48^. Consistently, neural representations in early visual areas are minimally influenced by prior knowledge, whereas DMN regions are significantly swayed by it ^49^. In addition, regions with more similar features tend to support parallel functions. For example, while FPCN and DMN generally exhibit negative functional connectivity, they share some similar attributes including long timescales, suggesting they can process inputs over longer periods. As a result, their neural representations are shaped by prior knowledge ^49^ and goal states maintained over time ^50^. These observations suggest that feature similarity can provide valuable information about the functional similarity of networks beyond functional connectivity. This unbiased, hypothesis-free analysis that combines machine learning with feature extraction can objectively pinpoint the specific subnetworks within DAN and DMN that demonstrate a heightened functional similarity to FPCN subnetworks and elucidate the functions of FPCN subnetworks.

We tested the hypothesis that FPCN-A and B parcels would be misclassified as belonging to different networks, reflecting their closest neighbors on the topographical spectrum ^23,44,48^: i.e., the classifier might misclassify parcels of FPCN-A as DAN and FPCN-B as DMN. We found classification accuracy was significantly greater than chance for each participant on each task (Fig. S2). Fig. 4 shows the top four networks with the highest prediction probabilities within the normalized confusion matrix, plus an additional comparator network; the probabilities for other networks were similar to chance level (Fig. S3).

By analyzing the classification output (i.e., the confusion matrix, Fig. S2), we found that FPCN-A and FPCN-B showed similarity to different networks. Specifically, FPCN-A was most likely to be misclassified as DAN-A and FPCN-B, while FPCN-B was most likely to be misclassified as FPCN-A and DMN-B (Fig. 4, Fig. S3). When the target network was FPCN-A, there was a higher probability that parcels would be misclassified as DAN-A than as DMN-B across rest and all the tasks (p < 0.05, FDR corrected; Fig. 4, Fig. S3, Table S1). Conversely, when the target network was FPCN-B, parcels were more likely to be misclassified as DMN-B than as DAN-A (p < 0.05, FDR corrected; Fig. 4, Fig S3, Table S1). These results indicate that the time-series characteristics of FPCN-A were more similar to DAN-A and those of FPCN-B were more similar to DMN-B. Therefore, subsequent analyses were focused on the functional similarity between DAN-A, DMN-B and the control subnetworks.

**Fig. 4.**
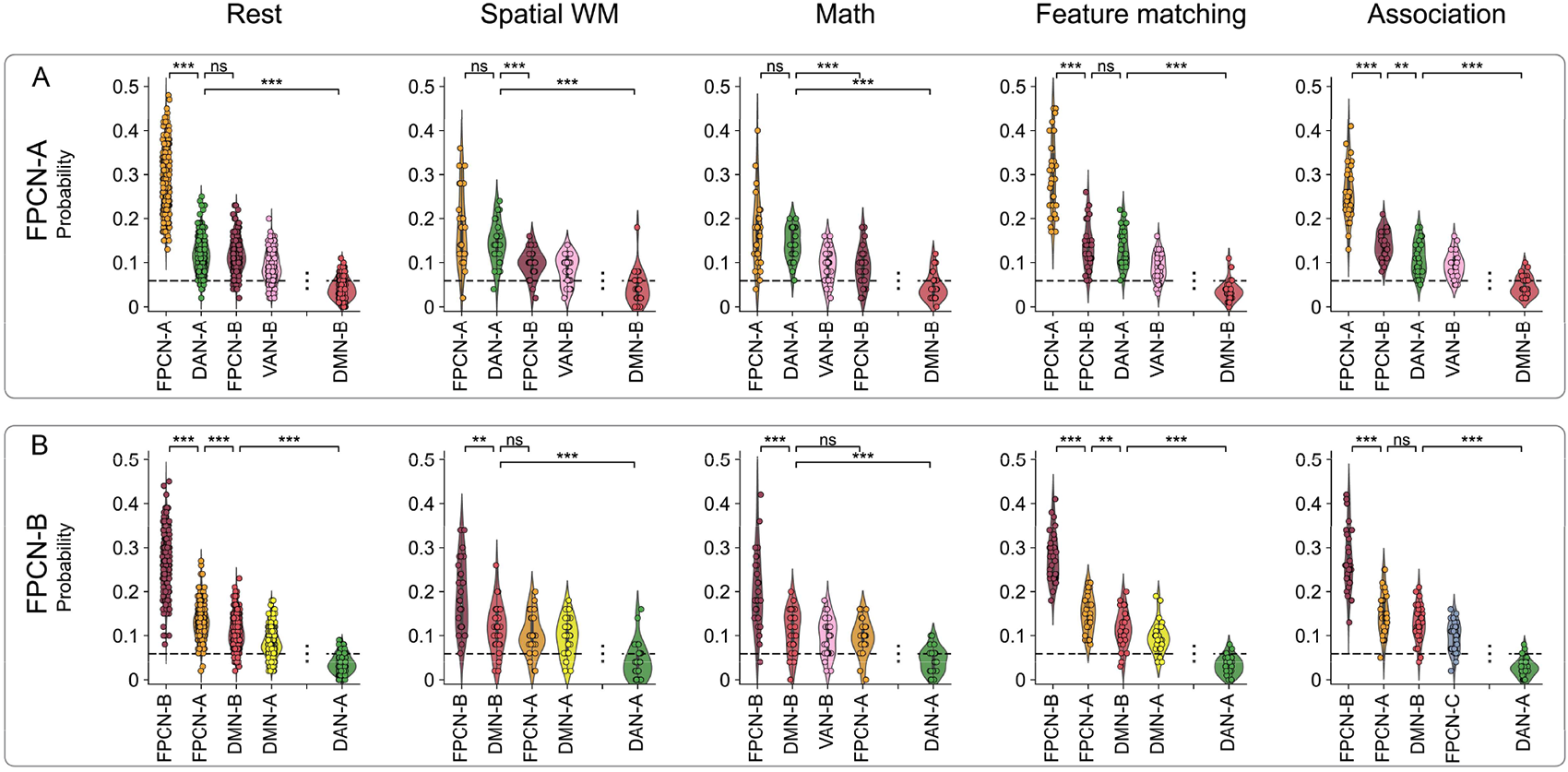
The probabilities of classifying FPCN-A and FPCN-B as each network (figure shows top four networks plus an additional comparator network). A: FPCN-A was most likely to be incorrectly classified as DAN-A at rest and in non-semantic tasks, and as FPCN-B in semantic tasks. It was also misclassified as VAN-B, but rarely misclassified as DMN-B. B: FPCN-B was most likely to be misclassified as DMN-B in non-semantic tasks and as FPCN-A in semantic tasks and at rest. It was also misclassified as DMN-A, and but rarely misclassified as DAN-A. Dashed lines represent the chance level.

#### 2.2.2 FPCN-A showed greater functional interaction with DAN-A, while FPCN-B showed greater coupling with DMN-B

To further characterise how topography affects functional interaction, we examined: (i) redundancy, which quantifies how much information about the brain’s future trajectory is shared across brain regions and (ii) functional connectivity between parcels, estimated by computing Pearson correlation coefficients between their time series. These measures revealed the same pattern as the network classification analysis: FPCN-A showed greater functional interaction with DAN-A, and FPCN-B showed greater functional interaction with DMN-B. These patterns were observed during rest and all the tasks (p < 0.05, FDR corrected; Fig. 5, Fig. S4, S5, S6, S8 and S9; Table S2, S3). In summary, both redundancy and connectivity analyses indicated greater functional similarity between control networks and the networks they were closer to (DAN; DMN); in this way, the topography of control subnetworks partly determines their functional tuning.

**Fig. 5.**
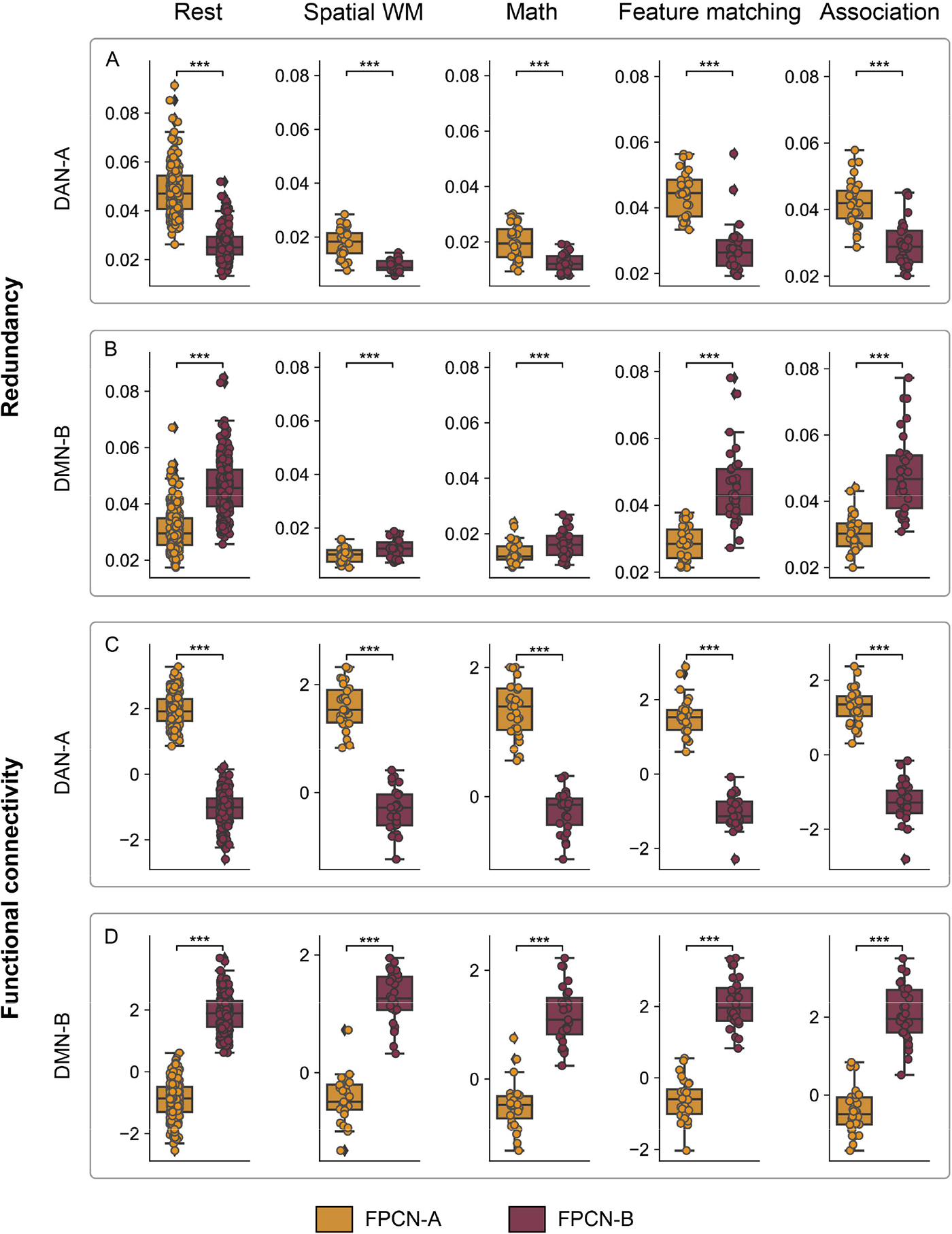
FPCN-A and FPCN-B coupled with different networks. FPCN-A always coupled with DAN-A while FPCN-B always coupled with DMN-B across different conditions. A and C: FPCN-A and DAN-A showed stronger redundancy and functional connectivity than FPCN-B and DAN-A. B and D: FPCN-B and DMN-B showed stronger redundancy and functional connectivity than FPCN-A and DMN-B.

#### 2.2.3 FPCN-A regions showed greater activation in more demanding conditions, while FPCN-B regions showed the opposite pattern

Given that FPCN-A and FPCN-B occupy different topographical positions on the cortical surface and show different strength of coupling with DAN-A and DMN-B, we might expect them to show different responses to control demands across tasks that differentially rely on dorsal attention and default networks. To test this hypothesis, we examined univariate effects of task difficulty at the whole brain level: in the working memory tasks, we contrasted hard and easy conditions, while the long-term memory tasks were designed to examine parametric effects of difficulty (manipulations of feature similarity in the semantic feature matching task and association strength in the semantic association task (see 4.3.3 and 4.3.4 for detailed information). All p-values were FDR corrected at p < 0.05. For clarity, we focus on the results of the four networks of interest below, and for completeness, the results of all the networks are shown in Fig. S7.

FPCN-A showed stronger activation in hard than easy trials across semantic and non-semantic tasks (Fig. 6). DAN-A also showed positive effects of difficulty in the spatial working memory, math and semantic feature matching tasks, yet showed deactivation during more difficult decisions in the semantic association task. In this way, FPCN-A showed a consistent response to difficulty irrespective of task context (as expected for regions of the “multiple demand network” ^12^, while DAN-A was not always engaged by semantic difficulty. In contrast, FPCN-B and DMN-B showed little or no positive response to difficulty, although FPCN-B parcels in dorsomedial prefrontal cortex did show a stronger response to more demanding decisions across math, feature matching and semantic association. FPCN-B typically deactivated in response to difficulty together with DMN-B (although this pattern was not observed for the semantic association task). Mean activation and deactivation patterns largely resembled effects of difficulty (SM 1.4 and Fig. S7). These results highlight a need to explain how topography not only supports differentiation of function within control subnetworks but also their flexible engagement in different task contexts.

**Fig. 6.**
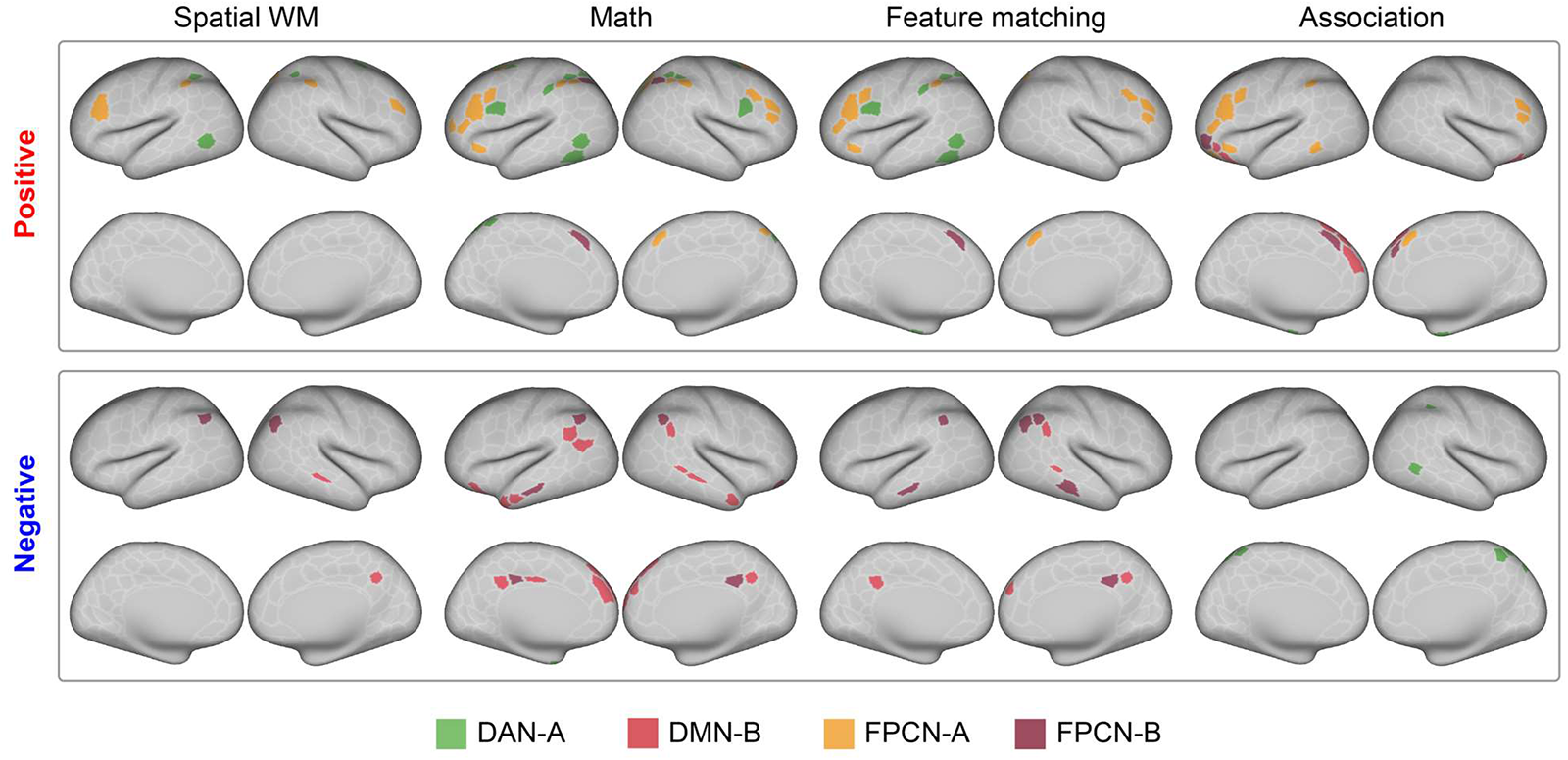
Modulation patterns of activation by task difficulty were similar for FPCN-A and DAN-A, and FPCN-B and DMN-B. FPCN-A showed stronger activation in hard than easy trials across semantic and non-semantic tasks. In contrast, FPCN-B typically deactivated in response to difficulty together with DMN-B. We examined univariate effects of task difficulty in the non-semantic tasks by contrasting hard and easy conditions. We examined the difficulty effect in the long-term memory tasks by examining parametric effects of feature similarity in the feature matching task and association strength in the association task, respectively.

### 2.3 The interaction patterns of FPCN-A varied across tasks, while FPCN-B showed stable patterns

Our final analyses tested how the interaction patterns of FPCN subnetworks varied across tasks tapping working and long-term memory. FPCN-A was recruited by difficult decisions across all four tasks, and was expected to show task dependent connectivity patterns, since domain-general control regions show a highly flexible pattern of coupling ^9,18^. However, regions that support controlled retrieval from long-term memory are thought not contribute to control across domains ^51–53^, and might show similar interaction patterns across tasks. Motivated by these findings, we asked whether FPCN-A would show greater interaction with DAN-A during non-semantic working memory tasks and with FPCN-B in semantic tasks (examining network misclassification, redundancy, and functional connectivity). We then asked whether FPCN-B would show consistent interaction with FPCN-A and DMN-B across task contexts. Features can differ depending on tasks and brain states ^54,55^, indicating their potential to depict variations in network coupling across tasks.

#### 2.3.1 FPCN-A was more likely to be misclassified as DAN-A in the non-semantic tasks but as FPCN-B in the semantic association task

We examined functional network similarity by analyzing the classification output (i.e., the confusion matrix) for each task, focussing on differences in the probability of misclassifying FPCN-A as DAN-A compared with FPCN-B across tasks. As expected, network similarity varied across tasks (Fig. 4A and Fig. S3). FPCN-A was more likely to be misclassified as DAN-A than as FPCN-B in non-semantic tasks (spatial working memory task: t = 3.913, p = 0.002; math task: t = 5.294, p < 0.001), but there was no difference between these networks in the feature matching task when participants needed to retrieve long-term memory according to external goals (t = 0.217, p = 0.83). FPCN-A was more likely to be misclassified as FPCN-B than as DAN-A in the semantic association task (t = -3.00, p = 0.008). All p values are FDR-corrected. In summary, FPCN-A varied its similarity to DAN-A and FPCN-B networks depending on the task state. This shows how misclassification effects go beyond patterns related to spatial adjacency to encompass task states. This shift in misclassification contradicts the typical assumption that networks and regions will always most closely resemble those that they are adjacent to on the cortical surface. While topography constrains the brain’s coupling pattern, it does not determine it entirely.

#### 2.3.2 FPCN-A showed greater functional interaction with DAN-A than with FPCN-B in the non-semantic tasks and the opposite pattern in the semantic tasks

Next, we considered differences in feature similarity, redundancy and functional connectivity between tasks, examining coupling of FPCN-A with DAN-A compared with FPCN-B. FPCN-A’s coupling to DAN-A compared with FPCN-B was always greater for the non-semantic than semantic tasks (p < 0.001; FWE corrected; Fig. 7). FPCN-A showed greater feature similarity, shared more information, and stronger functional connectivity with DAN-A in the spatial working memory and math tasks but coupled more with FPCN-B in the semantic tasks involving long-term memory (see SM 1.5 and 1.6. for detailed information for each task and each measure). Although the topography did not change, this shift in interaction challenges the prevailing assumption that networks and regions will consistently resemble their most adjacent cortical neighbours. The changing interaction patterns of FPCN-A provide convincing evidence to show that topography constrains but does not fully explain network connectivity.

#### 2.3.3 FPCN-B showed consistent interaction with FPCN-A and DMN-B across tasks

Having examined the flexible interaction patterns of FPCN-A, we asked if FPCN-B showed similar effects. We compared the probability that FPCN-B was misclassified as FPCN-A or as DMN-B in each task, as these were the most confusable networks for FPCN-B. As shown in Fig. 4B, misclassification was greater for FPCN-A than for DMN-B in the semantic feature matching task, when the participants needed to retrieve long-term memory according to external goals (t = 2.470, p = 0.019). FPCN-B was misclassified as FPCN-A and DMN-B equally often in the semantic association task (t = 1.811, p = 0.080) and the non-semantic tasks (spatial working memory task, t = -1.037, p = 0.309; math task, t = – 1.326, p = 0.245). All p values are FDR-corrected. These results show that although FPCN-B can resemble FPCN-A when people engage in controlled retrieval from long-term memory, this network also has a strong similarity with DMN-B across task contexts.

Finally, we quantified differences in feature similarity, redundancy and functional connectivity for FPCN-B across tasks, examining interaction of this network with DMN-B and FPCN-A. In contrast to FPCN-A’s flexibility, FPCN-B showed relatively stable interaction pattern. There were no differences in redundancy or functional connectivity across tasks (Fig. 7E and 6F; p > 0.05, uncorrected). FPCN-B showed equal redundancy with FPCN-A and DMN-B at rest and in all the tasks, with no differences between tasks (Fig. 7E, p > 0.05, uncorrected). FPCN-B always showed greater functional connectivity with DMN-B than with FPCN-A (Fig. 7F; p < 0.05, FWE corrected). FPCN-B showed greater feature similarity with FPCN-A than with DMN-B in the semantic feature matching task (t = 2.317, p = 0.033) but showed the opposite pattern in the spatial working memory task (t = -3.153, p = 0.005) and this task difference was significant (Fig. 7D; p < 0.002, FWE corrected). FPCN-B more closely resembled FPCN-A when people engaged in controlled retrieval from long-term memory but overall FPCN-B showed relatively consistent network interaction across tasks.

**Fig. 7.**
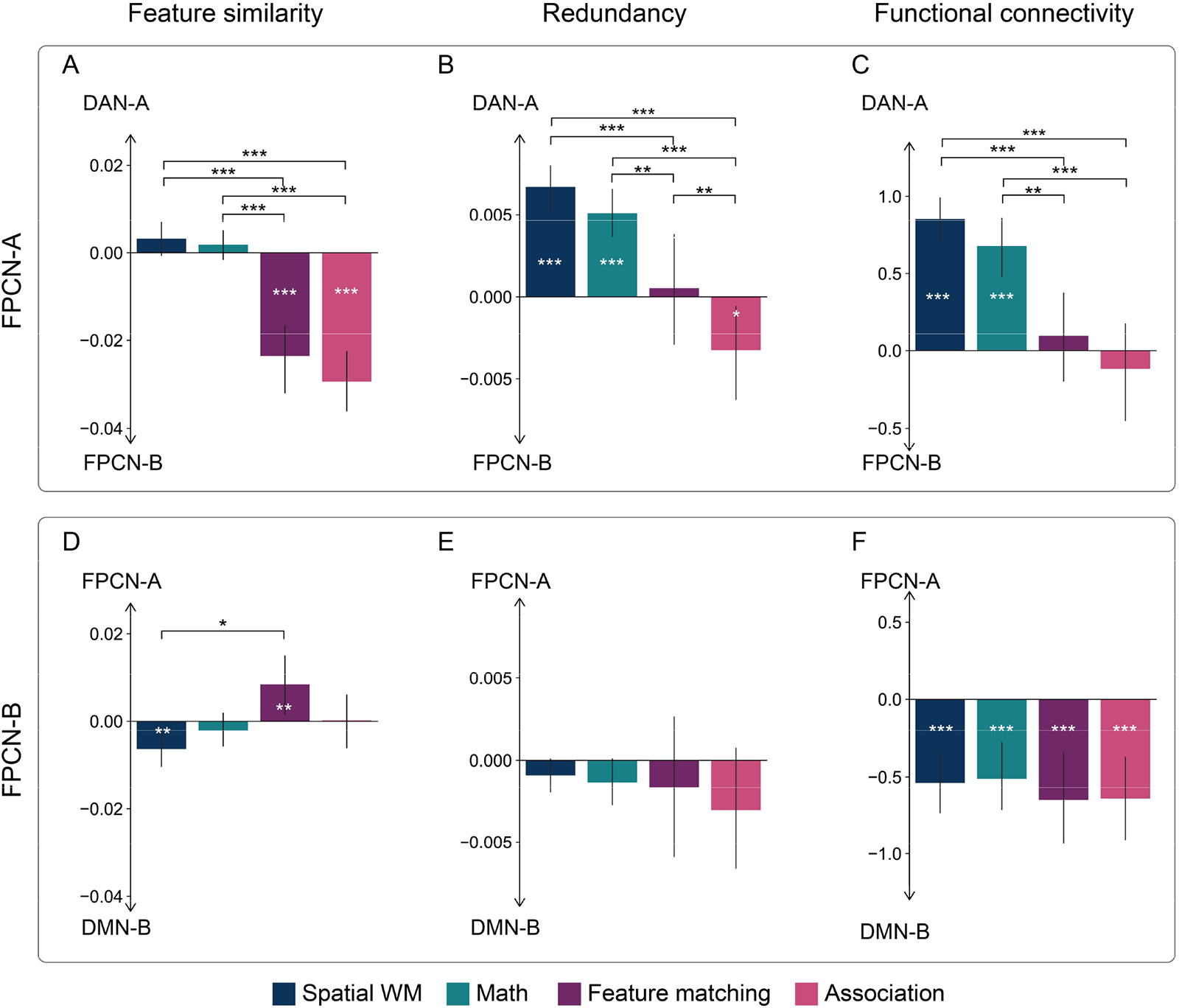
The interaction patterns of FPCN-A varied across tasks, while FPCN-B showed more stable patterns. The values in the top panel represent the interaction difference between FPCN-A and DAN-A versus FPCN-A and FPCN-B: positive values mean FPCN-A showed greater interaction with DAN-A than with FPCN-B. The values in the bottom panel represent the interaction difference between FPCN-B and FPCN-A versus FPCN-B and DMN-B: positive values mean FPCN-B showed greater interaction with FPCN-A than with DMN-B. A, B and C: The interaction differences in feature similarity, redundancy, and functional connectivity between FPCN-A and DAN-A versus FPCN-A and FPCN-B were greater in the non-semantic working memory tasks than in the semantic tasks. The task differences reflected greater interaction of FPCN-A with DAN-A than with FPCN-B in the non-semantic working memory tasks but the opposite pattern during the retrieval of knowledge from long-term memory. D, E, F: There were no interaction differences in redundancy, and functional connectivity between FPCN-B and FPCN-A versus FPCN-B and DMN-B across tasks. The difference in feature similarity was greater in the semantic feature matching task than in the spatial working memory task because FPCN-B showed greater interaction with FPCN-A than with DMN-B in the semantic feature matching task but showed the opposite pattern in the spatial working memory task. The white asterisk means the interaction difference was significant for that task at a specific threshold (*, p = 0.05; **, p = 0.01; ***, p = 0.001 by permutation based maximum t-tests, FWE corrected). The black asterisk means the interaction difference was significant across task pairs at a specific threshold (*, p = 0.05; **, p = 0.01; ***, p = 0.001 by permutation based maximum t-tests, FWE corrected).

## 3. Discussion

Our study shows that cortical topography, in conjunction with the underlying anatomy, provides a landscape in which flexible changes in neural function across situations can be understood. This landscape provides an architecture that supports the flexible deployment of distinct cognitive modes. By this account, FPCN subnetworks supporting distinct aspects of cognitive control are topographically positioned, and anatomically similar to their adjacent systems, allowing them to interact in a context-specific manner (Fig. 8). FPCN-A and FPCN-B are proximal to DAN and DMN respectively and this proximity is reflected by key anatomical features, including myelin content, the degree of cortical expansion, and cross-species similarity. This anatomical similarity complements the topographical organization of FPCN into adjacent subsystems and these aspects together account for important features of FPCN’s functional behavior. FPCN-A is more coupled with attention regions (particularly DAN-A), while FPCN-B is more coupled with memory regions (particularly DMN-B). Importantly, however, these relationships can change across contexts: FPCN-A shows stronger functional interaction to DAN-A during working memory and to FPCN-B during tasks more reliant on long-term memory. This shift in interaction challenges the prevailing assumption that networks and regions will consistently resemble their most adjacent cortical neighbors. FPCN-B shows more deactivation in response to demanding tasks and a pattern of interaction to both FPCN-A and DMN-B across tasks. The relative flexibility in FPCN-A and the relative stability of FPCN-B explains how people can utilize context-specific working memory and long-term memory to support flexible behavior.

These findings confirm that FPCN-A is closer to sensory-motor regions, positioned between DAN and FPCN-B on the cortical hierarchy, and can change its interaction to these networks depending on the task demands. In contrast, FPCN-B lies between FPCN-A and DMN-B and is coupled to both networks (Fig. 8). In this way, the topographical organisation of cognitive control regions allows FPCN-A to show the characteristics of a highly flexible ‘multiple-demand’ network, supporting executive control processes across domains ^12,13,56–61^, while the neighbouring FPCN-B shows the characteristics of a memory control network ^51,62–64^. This architecture might allow FPCN networks to be influenced by both DAN and DMN at different times, even though these networks are typically anti-correlated ^50,52,65^. In this way, our findings explain how previous studies have found both functional dissociations and similarities between FPCN and DMN (across different task contexts)^18,49,50,53^.

**Fig. 8.**
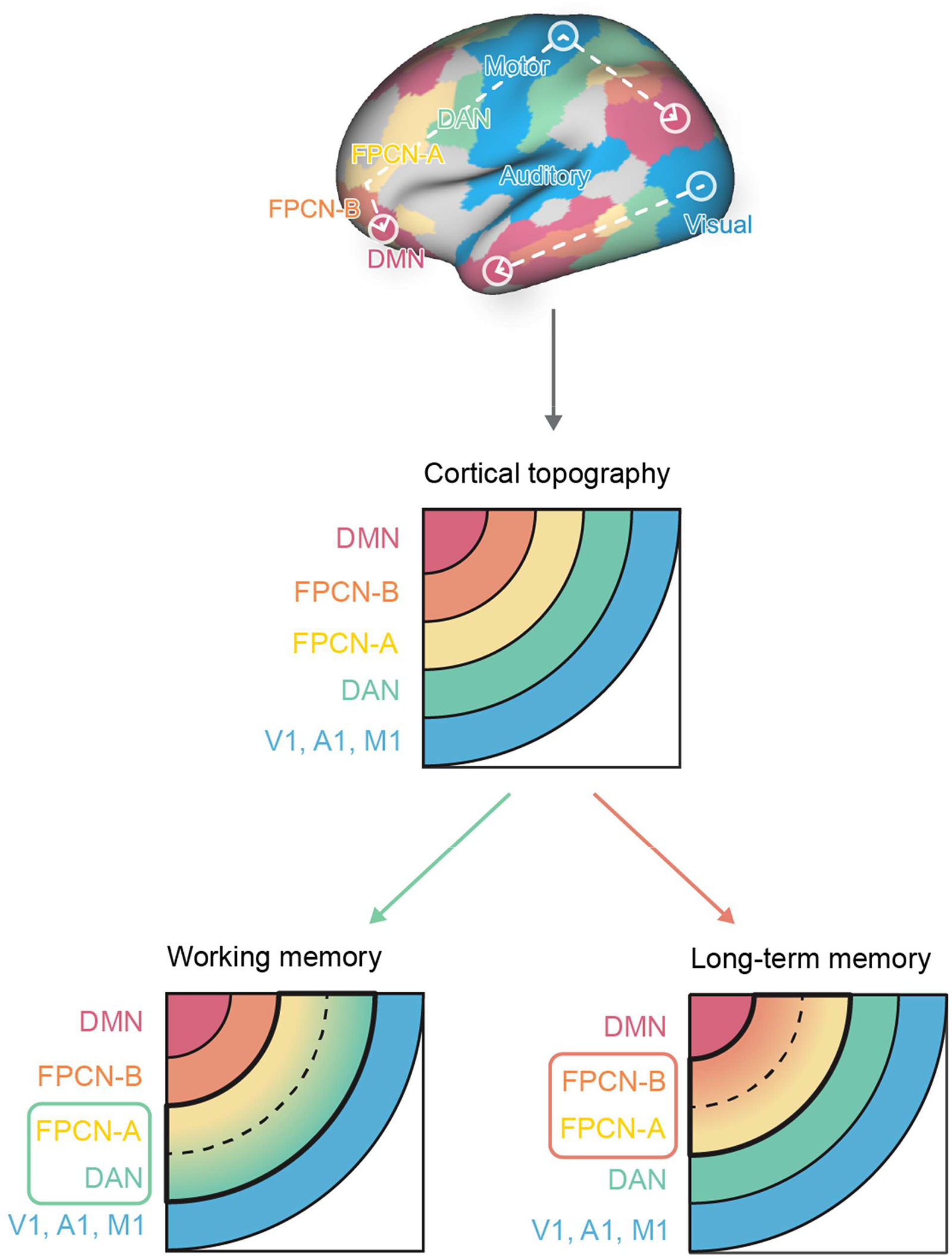
A schematic of the topographic relationships of networks and how this allows them to change their interaction patterns according to task demands to support flexible behavior, after Mesulam (1998). FPCN-A and FPCN-B are situated between DAN and DMN and they are influenced by their adjacent networks, allowing them to resolve competition and keep a functional balance between goal-oriented attentional mechanisms supporting working memory (drawing on DAN) and retrieval from long-term memory (supported by DMN). The topographic location of FPCN-A – situated between DAN and FPCN-B – allows it to shift its interaction towards attentional systems or memory systems according to task demands. In a changing environment, the engagement of externally oriented attentional mechanisms fractionates regions implicated in cognitive control, such that FPCN-A couples more with DAN-A than with FPCN-B in working memory task. This architecture might prevent interference from memory-based schema. When cognition instead needs to be guided by long-term memory, FPCN-A couples with FPCN-B, making this network less able to track fluctuating environmental changes but allowing it to integrate information from DMN.

The ability of FPCN-A to change its activity in a context specific manner indicates that this network might provide a ‘dynamic core’ controlling information flow by modulating network interactions in a context-sensitive fashion ^66^. Dynamic core regions are involved in multiple tasks, can integrate more specialized brain regions, and alter their baseline communication dynamics to support task-specific computations^67–70^. In a changing environment, interaction of FPCN-A with externally oriented attentional mechanisms appears to break apart the FPCN, such that FPCN-A couples more with DAN-A than with FPCN-B. This architecture might prevent interference from memory-based schema that conflict with the veridical state of the environment. When cognition instead needs to be guided by long-term memory, FPCN-A couples more with FPCN-B, making this network less able to track fluctuating environmental changes but allowing it to integrate information from DMN. Our findings are broadly consistent with the tethering hypothesis ^21^, which anticipates that proximity to sensory-motor regions constrains brain function. By extension, this account suggests that the relative separation of FPCN-A from DMN helps to protect information in working memory from biases that reflect expectations arising from conceptual knowledge and from fluctuations of emotion and motivation. Conversely, the proximity of FPCN-B to DMN allows it to support the retrieval of abstract and heteromodal concepts, and to reflect relevant information in memory even when this is at odds with the external environment ^71^. In tasks requiring controlled memory retrieval, these two control systems are expected to work together such that the representation of goals in working memory can interact with abstract semantic representations ^50,52,65^. Importantly, our study suggests that part of this flexibility may be reflected in the accompanying anatomical differences that mirror the topography of these networks.

There are of course limitations of the current study and remaining questions to be addressed. First, we are yet to establish whether the functional architecture we describe is relevant to all aspects of long-term memory, since our data are restricted to verbal semantic judgements. Future research should examine non-verbal semantic tasks and long-term memory beyond the semantic domain. Similarly, this architecture for flexible cognitive control may not extend to all working memory tasks, or to controlled perceptual-motor tasks. Other network interactions may also be important for flexible human cognition across a broader range of tasks, including the FPCN-C subnetwork (also described in the parcellation we used), which was not found to strongly contribute to the tasks in this study. Future studies can directly compare the flexibility of FPCN-A and FPCN-B using the same participants.

Future research should also assess the extent to which the mechanisms for flexible control described here relate to individual differences in cognitive performance and cognition in daily life, and whether topographical differences predict these individual differences in behaviour. A previous study that combined task performance and brain data using canonical correlation analysis found that working memory and long-term memory were two distinct neurocognitive modes that showed different associations with spontaneous thought in the lab ^72^. These modes might relate to the different patterns of interaction we identified between control networks; if so, individual differences in the relative strength or frequency of these controlled states might predict the extent to which people are able to control long-term memory retrieval and update working memory with goal-relevant information from the external world. They might also predict the kinds of thoughts that people are likely to have in their daily lives, with important implications for health, education, and wellbeing. Finally, while it is extremely challenging to examine the causal relationship between topography and function, computational simulations and studies of people with sensory impairment may help to elucidate this link.

In conclusion, this study shows how brain organisation supports mental flexibility. We establish that flexible human behavior is in part supported by the topographical separation of the frontoparietal system into subnetworks that are proximal to the dorsal attention network and default mode network. This topographical organization then allows distinct patterns of interaction to emerge in tasks underpinned by working memory and long-term knowledge, helping to explain how the brain maintains a functional balance between states that rely on top-down attention to the external environment compared with retrieval of information from long-term memory.

## 4. Materials and Methods

### 4.1. Experimental design

This study included three datasets, one open dataset – the Human Connectome Project (HCP) and two task fMRI datasets collected at the University of York, UK. We analysed the structural MRI (sMRI) and resting-state functional MRI (rsfMRI) data of 245 unrelated participants who completed all four resting-state scans from the S900 release of HCP dataset to investigate the cortical geometry, anatomical hierarchy, functional hierarchy, and functional connectivity patterns. To distinguish control processes related to recently presented information in working memory from control processes for long-term memory, we examined two non-semantic tasks requiring the maintenance of information in working memory which have been analysed before ^36,50,73^ and two semantic long-term memory tasks scanned in York, UK. In the non-semantic dataset, participants completed easy and hard spatial working memory and arithmetic tasks originally designed to localise domain general control regions ^13^. In the semantic dataset, participants performed a semantic feature matching task in which participants were asked to match probe and target concepts (presented as written words) according to color or shape and a semantic association task in which they were asked to decide if pairs of words were semantically associated or not. The contrast between non-semantic and semantic tasks allowed us to investigate how the interaction between control and other networks was modulated by task demands.

### 4.2. Participants

All participants were right-handed, native English speakers, with normal or corrected-to-normal vision and no history of psychiatric or neurological illness. All participants provided informed consent. For the HCP dataset, the study was approved by the Institutional Review Board of Washington University at St. Louis. For the University of York dataset, the research was approved by the York Neuroimaging Centre and Department of Psychology ethics committees.

The HCP sample involved data from 245 healthy volunteers (130 males, 115 females), aged 23 – 35 years (mean = 28.21, SD = 3.67) ^74^.

31 healthy adults (26 females; age: mean ± SD = 20.60 ± 1.68, range: 18 – 25 years) performed the spatial working memory and math tasks. One participant with incomplete data was removed. A functional run was excluded if (I) relative root mean square (RMS) framewise displacement was higher than 0.2 mm, (II) more than 15% of frames showed motion exceeding 0.25 mm, or (III) the accuracy of the behavior task was low (3SD below the mean). If only run of a task was left for a participant after exclusion, all their data for that task were removed. These exclusion criteria resulted in a final sample of 27 participants for both the spatial working memory task and the math task.

31 healthy adults performed the semantic long-term memory tasks (25 females; age: mean ± SD = 21.26 ± 2.93, range: 19 – 34 years). Using the same exclusion criteria above for the feature matching task, there were 23 participants with 4 runs, 4 participants with 3 runs, and 1 participant with 2 runs. For the association task, there were 24 participants with 4 runs, 3 participants with 3 runs, and 3 participants with 2 runs. Another 30 native English speakers who did not take part in the main fMRI experiment rated the color and shape similarity and semantic association strength for each word pair (21 females; age range: 18 – 24 years).

### 4.3. Task paradigms

#### 4.3.1. Spatial working memory task

Participants were required to maintain four or eight sequentially presented locations in a 3×4 grid ^75^, giving rise to easy and hard spatial working memory conditions. Stimuli were presented at the center of the screen across four steps. Each of these steps lasted for 1s and highlighted one location on the grid in the easy condition, and two locations in the hard condition. This was followed by a decision phase, which showed two grids side by side (i.e., two-alternative forced choice (2AFC) paradigm). One grid contained the locations shown on the previous four steps, while the other contained one or two locations in the wrong place. Participants indicated their response via a button press and feedback was immediately provided within in 2.5s. Each run consisted of 12 experimental blocks (6 blocks per condition and 4 trials in a 32 s block) and 4 fixation blocks (each 16 s long), resulting in a total time of 448 s.

#### 4.3.2. Math task

Participants were presented with an addition expression on the screen for 1.45s and, subsequently made a 2AFC decision indicating their solution within 1s. The easy condition used single-digit numbers while the hard condition used two-digit numbers. Each trial ended with a blank screen lasting for 0.1s. Each run consisted of 12 experimental blocks (with 4 trials per block) and 4 fixation blocks, resulting in a total time of 316s.

#### 4.3.3. Semantic feature matching task

Participants were required to make a yes/no decision matching probe and target concepts (presented as words) according to a particular semantic feature (color or shape), specified at the top of the screen during each trial. The feature prompt, probe word, and target words were presented simultaneously. Half of the trials were matching trials in which participants would be expected to identify shared features; while half of the trials were non-matching trials in which participants would not be expected to identify shared features. For example, in a color matching trial participants would answer ‘yes’ to the word-pair DALMATIANS – COWS, due to their color similarity, whereas they would answer ‘no’ to COAL -TOOTH as they do not share a similar color.

We parametrically manipulated the degree of feature similarity between the probe and target concepts, using semantic feature similarity ratings taken from a separate group of 30 participants on a 5-point Likert Scale. For instance, in color-matching trials, the degree of color similarity between DALMATIANS and COWS was found to be very high (i.e., 4.8), while that between PUMA and LION was relatively low (i.e., 4.0), despite the participants’ belief that the two trials had similar color. Conversely, in color non-matching trials, the degree of color similarity between CROW and HUMMINGBIRD was relatively high (i.e., 2.5), whereas that between COAL and TOOTH was very low (i.e., 1.2), even though the participants perceived no similarity in color. For the matching trials, greater feature similarity facilitates the decision-making process, while for the non-matching trials, greater feature similarity makes the decision more difficult. This parametric design allowed us to model the effect of the difficulty of semantic decision in the neural data, and test whether control subnetworks showed similar or opposite activation patterns.

This task included four runs and two conditions (two features: color and shape), presented in a mixed design. Each run consisted of four experimental blocks (two 2 min 30 s blocks per feature), resulting in a total time of 10 min 12 s. In each block, 20 trials were presented in a rapid event-related design. In order to maximize the statistical power of the rapid event-related fMRI data analysis, the stimuli were presented with a temporal jitter randomized from trial to trial ^76^. The inter-trial interval varied from 3 to 5 s. Each trial started with a fixation, followed by the feature, probe word, and target word presented centrally on the screen, triggering the onset of the decision-making period. The feature, probe word, and target word remained visible until the participant responded, or for a maximum of 3 s. The condition order was counterbalanced across runs and run order was counterbalanced across participants. Half of the participants pressed a button with their right index finger to indicate a matching trial and responded with their right middle finger to indicate a non-matching trial. Half of the participants pressed the opposite buttons.

#### 4.3.4. Semantic association task

Participants were asked to decide if pairs of words were semantically associated or not (i.e., yes/no decision as above) based on their own experience. Overall, there were roughly equal numbers of ‘related’ and ‘unrelated’ responses across participants. The same stimuli were used in the semantic feature matching task and semantic association task. For example, DALMATIANS and COWS are semantically related; COAL and TOOTH are not. The feature and association tasks were separated by one week. Similarly, we parametrically manipulated the semantic association strength between the probe and target concepts, using semantic association strength ratings taken from a separate group of 30 participants on a 5-point Likert Scale. For example, in related trials, the association strength between PUMA and LION is very strong while for COWS and WHALE it is relatively weak (although they are still both animals). In non-related trials, the association strength between KINGFISHER and SCORPION is relatively high while BANANA and BRICK is very low although participants thought neither were related. For the related trials, stronger associations would facilitate decision making, while for unrelated trials, stronger associations interfere with the decision making. This parametric design allowed us to model the effect of the difficulty of semantic decision in the neural data, and test whether control subnetworks showed similar or opposite activation patterns.

This task included four runs, presented in a rapid event-related design. Each run consisted of 80 trials, with about half being related and half being unrelated trials. The procedure was the same as the feature matching task except only two words were presented on the screen.

### 4.4. Image acquisition

#### 4.4.1. Image acquisition of HCP dataset

MRI acquisition protocols of the HCP dataset have been previously described ^74,77^. Images were acquired using a customized 3T Siemens Connectome scanner having a 100 mT/m SC72 gradient set and using a standard Siemens 32-channel radiofrequency receive head coil. Participants underwent the following scans: structural (at least one T1-weighted (T1w) MPRAGE and one 3D T2-weighted (T2w) SPACE scan at 0.7-mm isotropic resolution), rsfMRI (4 runs ×14 min and 33 s). Since not all participants completed all scans, we only included 339 unrelated participants from the S900 release. Whole-brain rsfMRI and task fMRI data were acquired using identical multi-band echo planar imaging (EPI) sequence parameters of 2-mm isotropic resolution with a TR = 720 ms.

Subjects were considered for data exclusion based on the mean and mean absolute deviation of the relative root-mean-square motion across either four rsfMRI scans or one dMRI scan, resulting in four summary motion measures. If a subject exceeded 1.5 times the interquartile range (in the adverse direction) of the measurement distribution in two or more of these measures, the subject was excluded. In addition, functional runs were flagged for exclusion if more than 25% of frames exceeded 0.2 mm frame-wise displacement (FD_power). These above exclusion criteria were established before performing the analysis ^78,79^. The data of 91 participants was excluded because of excessive head motion and the data of another 3 participants was excluded because their resting data did not have all the time points. In total, the data of 245 participants was analysed after exclusions.

#### 4.4.2. Image acquisition of York Non-semantic dataset

MRI acquisition protocols have been described previously ^50,73^. Structural and functional data were collected on a Siemens Prisma 3T MRI scanner at the York Neuroimaging Centre. The scanning protocols included a T1-weighted MPRAGE sequence with whole-brain coverage. The structural scan used: acquisition matrix of 176 × 256 × 256 and voxel size 1 × 1 × 1 mm^3^, repetition time (TR) = 2300 ms, and echo time (TE) = 2.26 ms. Functional data were acquired using an EPI sequence with an 800 flip angle and using GRAPPA with an acceleration factor of 2 in 3 x 3 x 4 mm voxels in 64-axial slices. The functional scan used: 55 3-mm-thick slices acquired in an interleaved order (with 33% distance factor), TR = 3000 ms, TE = 15 ms, FoV = 192 mm.

#### 4.4.3. Image acquisition of York Semantic dataset

Whole brain structural and functional MRI data were acquired using a 3T Siemens MRI scanner utilising a 64-channel head coil, tuned to 123 MHz at York Neuroimaging Centre, University of York. The functional runs were acquired using a multi-band multi-echo (MBME) EPI sequence, each 11.45 minutes long (TR=1.5 s; TE = 12, 24.83, 37.66 ms; 48 interleaved slices per volume with slice thickness of 3 mm (no slice gap); FoV = 24 cm (resolution matrix = 3x3x3; 80x80); 75° flip angle; 455 volumes per run; 7/8 partial Fourier encoding and GRAPPA (acceleration factor = 3, 36 ref. lines); multi-band acceleration factor = 2). Structural T1-weighted images were acquired using an MPRAGE sequence (TR = 2.3 s, TE = 2.3 s; voxel size = 1x1x1 isotropic; 176 slices; flip angle = 8°; FoV= 256 mm; interleaved slice ordering). We also collected a high-resolution T2-weighted (T2w) scan using an echo-planar imaging sequence (TR = 3.2 s, TE = 56 ms, flip angle = 120°; 176 slices, voxel size = 1x1x1 isotropic; Fov = 256 mm).

### 4.5. Image pre-processing

#### 4.5.1. Image pre-processing of HCP dataset

We used HCP’s minimal pre-processing pipelines ^74^. Briefly, for each subject, structural images (T1w and T2w) were corrected for spatial distortions. FreeSurfer v5.3 was used for accurate extraction of cortical surfaces and segmentation of subcortical structures ^80,81^. To align subcortical structures across subjects, structural images were registered using non-linear volume registration to the Montreal Neurological Institute (MNI152) space. Functional images (rest and task) were corrected for spatial distortions, head motion, and mapped from volume to surface space using ribbon-constrained volume to surface mapping.

Subcortical data were also projected to the set of extracted subcortical structure voxels and combined with the surface data to form the standard CIFTI grayordinate space. Data were smoothed by a 2-mm FWHM kernel in the grayordinates space. Rest data were additionally identically cleaned for spatially specific noise using spatial ICA+FIX ^82^ and global structured noise using temporal ICA ^83^. For accurate cross-subject registration of cortical surfaces, a multimodal surface matching (MSM) algorithm ^84^ was used to optimize the alignment of cortical areas based on features from different modalities. MSMSulc (“sulc”: cortical folds average convexity) was used to initialize MSMAll, which then utilized myelin, resting-state network, and rfMRI visuotopic maps. Myelin maps were computed using the ratio of T1w/T2w images ^82^. The HCP’s minimally preprocessed data include cortical thickness maps (generated based on the standardized FreeSurfer pipeline with combined T1-/T2-reconstruction). For this study, the standard-resolution cortical thickness maps (32k mesh) were used.

#### 4.5.2. Image pre-processing of York Non-semantic and Semantic dataset

The York datasets were preprocessed using fMRIPrep 20.2.1 [^85^, RRID:SCR_016216], which is based on Nipype 1.5.1 [^86^, RRID:SCR_002502].

##### 4.5.2.1. Anatomical data preprocessing

The T1w image was corrected for intensity non-uniformity (INU) with N4BiasFieldCorrection ^87^, distributed with ANTs 2.3.3 [^88^, RRID:SCR_004757], and used as T1w-reference throughout the workflow. The T1w-reference was then skull-stripped with a Nipype implementation of the antsBrainExtraction.sh workflow (from ANTs), using OASIS30ANTs as target template. Brain tissue segmentation of cerebrospinal fluid (CSF), white-matter (WM) and gray-matter (GM) was performed on the brain-extracted T1w using fast FSL 5.0.9 [^89^, RRID:SCR_002823]. Brain surfaces were reconstructed using recon-all from FreeSurfer 6.0.1 [^90^, RRID:SCR_001847], and the brain mask estimated previously was refined with a custom variation of the method to reconcile ANTs-derived and FreeSurfer-derived segmentations of the cortical gray-matter of Mindboggle [^91^, RRID:SCR_002438]. Volume-based spatial normalization to two standard spaces (MNI152NLin2009cAsym, MNI152NLin6Asym) was performed through nonlinear registration with antsRegistration (ANTs 2.3.3), using brain-extracted versions of both T1w reference and the T1w template. The following templates were selected for spatial normalization: ICBM 152 Nonlinear Asymmetrical template version 2009c [^92^, RRID:SCR_008796; TemplateFlow ID: MNI152NLin2009cAsym], FSL’s MNI ICBM 152 non-linear 6th Generation Asymmetric Average Brain Stereotaxic Registration Model [^93^, RRID:SCR_002823; TemplateFlow ID: MNI152NLin6Asym].

##### 4.5.5.2. Functional data preprocessing

For each of the BOLD runs per subject, the following preprocessing was performed. First, a reference volume and its skull-stripped version were generated using a custom methodology of fMRIPrep. A B0-nonuniformity map (or fieldmap) was estimated based on a phase-difference map calculated with a dual-echo GRE (gradient-recall echo) sequence, processed with a custom workflow of SDCFlows inspired by the epidewarp.fsl script and further improvements in HCP Pipelines ^74^. The fieldmap was then co-registered to the target EPI reference run and converted to a displacements field map (amenable to registration tools such as ANTs) with FSL’s fugue and other SDCflows tools. Based on the estimated susceptibility distortion, a corrected EPI reference was calculated for a more accurate co-registration with the anatomical reference. The BOLD reference was then co-registered to the T1w reference using bbregister (FreeSurfer) which implements boundary-based registration ^94^. Co-registration was configured with six degrees of freedom. Head-motion parameters with respect to the BOLD reference (transformation matrices, and six corresponding rotation and translation parameters) were estimated before any spatiotemporal filtering using mcflirt (FSL 5.0.9,^95^). BOLD runs were slice-time corrected using 3dTshift from AFNI 20160207 [(27), RRID:SCR_005927]. Since multi-echo BOLD data was supplied in the York Semantic dataset, the tedana T2* workflow was used to generate an adaptive T2* map and optimally weighted combination of all supplied single echo time series. This optimally combined time series was then carried forward for all subsequent preprocessing steps. The BOLD time-series were resampled onto the following surfaces (FreeSurfer reconstruction nomenclature): fsaverage. Grayordinates files ^74^ containing 91k samples were also generated using the highest-resolution fsaverage as intermediate standardized surface space. Several confounding time-series were calculated based on the preprocessed BOLD: framewise displacement (FD), DVARS and three region-wise global signals. FD was computed using two formulations following previous work (absolute sum of relative motion; ^97^, relative root mean square displacement between affines ^95^). FD and DVARS were calculated for each functional run, both using their implementations in Nipype. ^97^. Three global signals were extracted within the CSF, the WM, and the whole-brain masks. The confound time series derived from head motion estimates and global signals were expanded with the inclusion of temporal derivatives and quadratic terms for each ^98^. Frames that exceeded a threshold of 0.5 mm FD or 1.5 standardized DVARS were annotated as motion outliers. All resamplings were performed with a single interpolation step by composing all the pertinent transformations (i.e., head-motion transform matrices, susceptibility distortion correction when available, and co-registrations to anatomical and output spaces). Gridded (volumetric) resamplings were performed using antsApplyTransforms (ANTs), configured with Lanczos interpolation to minimize the smoothing effects of other kernels ^99^. Non-gridded (surface) resamplings were performed using mri_vol2surf (FreeSurfer). fMRIPrep used Nilearn 0.6.2 [^100^ RRID:SCR_001362], mostly within the functional processing workflow. The resulting data were in CIFTI 64k-vertex grayordinate space. The left hemisphere had 29696 vertices and right hemisphere had 29716 vertices in total after removing the medial wall.

Post-processing of the outputs of fMRIPrep version 20.2.1 ^85^ was performed using the eXtensible Connectivity Pipeline (XCP) ^98,101^. For each CIFTI run per subject, the following post-processing was performed: before nuisance regression and filtering any volumes with framewise-displacement greater than 0.3 mm ^97,98^ were flagged as outliers and excluded from nuisance regression. In total, 36 nuisance regressors were selected from the nuisance confound matrices of fMRIPrep output. These nuisance regressors included six motion parameters, global signal, mean white matter, and mean CSF signal with their temporal derivatives, and the quadratic expansion of six motion parameters, tissue signals and their temporal derivatives ^98,101,102^. These nuisance variables were accounted for in the BOLD data using linear regression – as implemented in Scikit-Learn 0.24.2 ^103^. Residual timeseries from this regression were then band-pass filtered to retain signals within the 0.01-0.08 Hz frequency band. The processed BOLD was smoothed using Connectome Workbench with a gaussian kernel size of 6.0 mm (FWHM). Processed functional timeseries were extracted from residual BOLD using Connectome Workbench ^74^ . Many internal operations of XCP use Nibabel ^100^, numpy ^104^, and scipy ^104^.

### 4.6. Structural and task fMRI analysis

#### 4.6.1. Individual-specific parcellation

Considering the anatomical and functional variability across individuals ^105–108^, we estimated individual-specific areal-level parcellation using a multi-session hierarchical Bayesian model (MS-HBM) ^30,109^. To estimate individual-specific parcellation, we acquired ‘‘pseudo-resting state’’ timeseries in which the task activation model was regressed from feature matching, semantic association, spatial working memory, and math fMRI data ^110^ using xcp_d (https://github.com/PennLINC/xcp_d). The task activation model and nuisance matrix were regressed out using AFNI’s3dTproject (for similar implementation, see Cui et al.^111^_)._

Using a group atlas, this method calculates inter-subject resting-state functional connectivity variability, intra-subject resting-state functional connectivity variability, and finally parcellates for each single subject based on this prior information. As in Kong et al. ^30,109^, we used MS-HBM to define 400 individualized parcels belonging to 17 discrete individualized networks for each participant in which control network was divided into 3 sub-networks, allowing us to explore the heterogeneity of control network. Specifically, we calculated all participants’ connectivity profiles, created the group parcellation using the average connectivity profile of all participants, estimated the inter-subject and intra-subject connectivity variability, and finally calculated each participant’s individualized parcellation. This parcellation imposed the Markov random filed (MRF) spatial prior. We used a well-known areal-level parcellation approach, i.e., the local gradient approach (gMS-HBM), which detects local abrupt changes (i.e., gradients) in resting-state functional connectivity across the cortex ^112^. A previous study ^113^ has suggested combining local gradient ^33,112^ and global clustering ^31^ approaches for estimating areal-level parcellations. Therefore, we complemented the spatial contiguity prior in contiguous MS-HBM (cMS-HBM) with a prior based on local gradients in resting-state functional connectivity, which encouraged adjacent brain locations with gentle changes in functional connectivity to be grouped into the same parcel. We used the pair of parameters (i.e., beta value = 50, w = 30 and c = 30), which was optimized using our own dataset. The same parameters were also used in Kong et al. ^30^. Vertices were parcellated into 400 cortical regions (200 per hemisphere). To parcellate each of these parcels, we calculated the average time series of enclosed vertices to get better signal noise ratio (SNR) using Connectome Workbench software. This parcel-based time series was used for all the following analyses. The same method and parameters were used to generate the individual-specific parcellation for the participants in the HCP dataset using the resting-state time series except that the task regression was not performed.

##### 4.6.1.1 Homogeneity of parcels

To evaluate whether a functional parcellation is successful, parcel homogeneity is commonly used ^30,33,109^. Parcel homogeneity was calculated as the average Pearson’s correlations between fMRI time courses of all pairs of vertices within each parcel, adjusted for parcel size and summed across parcels ^30,109,113^. Higher homogeneity means that vertices within the same parcel share more similar time courses and indicates better parcellation quality. To summarize the parcel homogeneity, we averaged the homogeneity value across parcels. We calculated the parcel homogeneity for each run of each participant for each task using the individual-specific parcellation and then average them across runs for each participant for each task. We also calculated the parcel homogeneity using canonical Yeo 17-network group atlas. Using the resting state data of the HCP dataset, Kong et al ^30^ have demonstrated that homogeneity within MS-HBM-based individualized parcels was greater than that in the canonical Yeo 17-network group atlas that does not consider variation in functional neuroanatomy. The similar pattern was observed using York Non-semantic and Semantic datasets (Fig. S1). We also observed that the homogeneity of the semantic tasks of York Semantic dataset were higher than the non-semantic tasks of York Non-semantic dataset. The potential reason might be that we collected T2-weighted images for the semantic tasks to improve the skull-stripping, giving a better outcome for pial surface reconstruction. Given the known heterogeneity within the FPCN, as well as within the DAN and DMN, we did not merge any subnetworks as was done by Dixon et al. (2018) and Murphy et al. (2020).

#### 4.6.2. Cortical geometry – global minimum distance to primary sensory-motor landmarks

To reveal how the physical proximity to structural landmarks corresponding to primary sensory-motor regions influences the function of regions, we calculated the geodesic distance between each parcel and key landmarks associated with primary visual, auditory and somatomotor cortex. We used these values to identify the minimum geodesic distance to primary sensory-motor regions for each parcel. Three landmarks were used: central sulcus, which is the topographical landmark corresponding to primary somatosensory/motor cortex; temporal transverse sulcus, which provides a landmark for primary auditory cortex; and calcarine sulcus, marking the location of primary visual cortex. Since cortical folding patterns vary across participants, and individual variability in cortical folding increases with cortical surface area, both the shapes of these landmarks and the number of vertices within each landmark might show individual variability ^114^. We used participant-specific landmark label files to locate the participant-specific vertices belonging to each landmark and participant-specific parcellation to locate the vertices within each parcel ^36^.

Geodesic distance along the “midthickness” of the cortical surface (halfway between the pial and white matter) was calculated using the Connectome Workbench software with an algorithm that measures the shortest path between two vertices on a triangular surface mesh ^115,116^. This method returns distance values that are independent of mesh density. Geodesic distance was extracted from surface geometry (GIFTI) files, following surface-based registration ^84^. To ensure that the shortest paths would only pass through the cortex, vertices representing the medial wall were removed from the triangular mesh in this analysis.

We calculated the minimum geodesic distance between each vertex and each landmark. Specifically, for the landmark central sulcus, we calculated the geodesic distance between vertex *i* outside central sulcus and each vertex within the central sulcus (defined for each individual; see above). We then found vertex *j* within the central sulcus closest to vertex *i*, and extracted this value as the minimum geodesic distance between vertex *i* and this landmark. To compute the minimum geodesic distance between parcel *k* and the central sulcus, we computed the average minimum distance across all the grayordinate vertices in parcel *k* to the vertices within the central sulcus. We used the same procedure to calculate minimum geodesic distance between each parcel and all three sensory-motor landmarks (central sulcus and temporal transverse sulci as well as calcarine sulcus). From these three minimum geodesic distances identified between parcel *k* and sensory-motor landmarks, we then selected the lowest distance value (i.e., the landmark that was closest to parcel *k*) to define the global minimum distance to sensory-motor regions for parcel *k*. Then we averaged the mean minimum distance of all the parcels within each network for each participant and then sorted the networks by the mean minimum distance across participants. Finally, we examined whether mean minimum distance of FPCN-A and FPCN-B were different by performing a paired t-test.

#### 4.6.3. Anatomical hierarchy – myelin content and cortical thickness

We measured myelin which is a non-invasive and valid proxy for anatomical hierarchy and captures the anatomical hierarchy better than cortical thickness ^22^. The gray-matter myelin content can be measured via the cortical T1w/T2w map, which is a structural neuroimaging map defined by the contrast ratio of T1- to T2-weighted (T1w/T2w) magnetic resonance images ^24,35,74,82^.

Human T1w/T2w maps were obtained from the HCP in the surface-based CIFTI file format. To produce the T1w/T2w maps, high resolution T1- and T2-weighted images were first registered to a standard reference space using a state-of-the-art areal-feature-based technique ^24,35,74^.

Each participant has a myelin map, with each vertex having a myelin value (i.e., T1w/T2w). We calculated the myelin value of each parcel by averaging the myelin values of all the vertices within this parcel for each participant using the individual-specific parcellation. Similarly, we calculated the myelin value of each network by averaging the myelin values of all the parcels within each network for each participant and then sorted the networks by the mean myelin value across participants. Finally, we examined whether the myelin values of the FPCN-A and FPCN-B were different by performing paired t-tests.

Cortical thickness coarsely tracks changes in cytoarchitecture and myelin content, and can be viewed as a pragmatic surrogate for cortical microstructure. Therefore, we also examined cortical thickness of each network which measures the width of gray matter of cortex across networks. The procedure was as above, except we used the cortical thickness maps and extracted the cortical thickness of each vertex of each participant.

#### 4.6.4. Function hierarchy – principal connectivity gradient analysis

To examine the relative position of networks on the principal gradient axis of intrinsic connectivity, we performed dimension reduction analysis on resting state functional connectivity matrix of HCP dataset. First, the resting-state functional connectivity for each run of each participant was calculated using the method in 4.6.10. These individual connectivity matrices were then averaged to calculate a group-averaged connectivity matrix. The Brainspace Toolbox ^117^ was used to extract ten group-level gradients from the group-averaged connectivity matrix (dimension reduction technique = diffusion embedding, kernel = None, sparsity = 0.9), in line with previous studies ^118^. Using identical parameters, gradients were then calculated for each individual using their average 400 × 400 resting state functional connectivity matrix across four runs. These individual-level gradient maps were aligned to the group-level gradient maps using Procrustes rotation to improve comparison between the group-level gradients and individual-level gradients (N iterations = 10). This analysis resulted in ten group-level gradients and ten individual-level gradients for each participant explaining maximal whole-brain connectivity variance in descending order. For 238 out of 245 participants, the first gradient explaining maximal variance, was the principal gradient which captures the separation between unimodal and transmodal regions ^20^. Next, we averaged the first gradient values of all the parcels within each network for each participant whose first gradient was the principal gradient and then sorted the networks by the mean gradient values across participants. Finally, we examined whether the gradient values of FPCN-A and FPCN-B were different by performing paired t-tests.

#### 4.6.5. Comparing the evolutionary expansion and cross-species similarity between FPCN-A and FPCN-B

We used the evolutionary expansion map and cross-species similarity map ^29^ from https://github.com/TingsterX/alignment_macaque-human. Surface areal expansion map was calculated as the human area divided by macaque area at each of corresponding vertex on human and macaque surfaces ^29^. Cross-species similarity is calculated by comparing the similarity of whole brain patterns of functional connectivity in macaques and humans ^29^. We extracted the value of each vertex and calculated the mean value of each parcel of the group parcellation. Finally, we compared the value between FPCN-A and FPCN-B by conducting independent t-tests.

#### 4.6.6. Feature extraction of the time series data

We used the highly comparative time-series analysis toolbox, hctsa^119,120^, to extract massive features of the time-series. Using a time-series dataset, hctsa allowed us to transform each timeseries to a set of over 7,700 features that each encodes a different scientific analysis method ^119,120^. The extracted features include, but are not limited to, distributional properties, entropy and variability, autocorrelation, time-delay embeddings, and nonlinear properties of a given time-series ^120,121^. The hctsa feature extraction analysis was performed on the parcellated fMRI time-series of each participant, each task and each run separately. Following the feature extraction procedure, the outputs of the operations that produced errors were removed and the remaining features (about 6900 features) were normalized across parcels using an outlier-robust sigmoidal transform. The normalized feature matrix (400 parcels × about 7000 features × 4 runs) was used for further classification analysis and feature similarity analysis.

#### 4.6.7. Classification analysis

To reveal the network similarity in a data-driven approach, we performed a classification analysis using the normalized feature matrix.

##### 4.6.7.1. Balanced accuracy of classification analysis

To examine whether we could classify the network labels of each parcel using the extracted massive features, we performed a classification task to investigate how accurately a classifier can learn a mapping from time-series features of parcels to network labels of these parcels. We combined the normalized features of all the runs of each task for each participant and then performed the multi-class classification (17 networks labels) using scikit-learn ^103^, which is a machine learning library written in Python.

For the multi-class classification, we trained linear support vector machine classifiers (sklearn.svm.SVC) to find the hyperplane that maximally separates the samples belonging to the difference classes. The parcel numbers of networks are different. For example, both FPCN-A and FPCN-B have 25 parcels, while Control-C has 23 parcels. Due to imbalance of observations across the networks, we reported balanced classification accuracy^122,123^. Specifically, the balanced accuracy was calculated as the arithmetic mean of sensitivity, (i.e., true positive rate which measures the proportion of real positives that are correctly predicted out of total positive prediction made by the model), and specificity, (i.e., true negative rate which measures the proportion of correctly identified negatives over the total negative prediction made by the model). We performed 5-fold cross-validation to prevent overfitting leading to optimistic performance estimates. To examine whether the classification accuracy was significantly greater than the chance level for each participant and each task, we performed the permutation-based multi-class classification analysis in which we randomly shuffled the network labels 1000 times within all the runs for each participant for each task. This established an empirical distribution of classification accuracy scores under the null hypothesis where there is no association between features and network labels ^124^.

The data used for classification have about 7000 features and 1600 samples (400 parcels × 4 runs) in the semantic tasks and 800 samples (400 samples × 2 runs) in the non-semantic tasks. Machine learning with many more features than samples is challenging, due to the so-called curse of dimensionality. The curse of dimensionality describes the explosive nature of increasing data dimensions; this increase in the dimensions might increase the noise and redundancy during its analysis. It has been shown that with a fixed number of training samples, the predictive power of any classifier first increases as the number of dimensions increase, but after a certain value of number of dimensions, the performance deteriorates ^125^. To explore the curse of dimensionality, we performed the classification using all the features and part of the features, respectively. Parts of the features were the top features that make major contributions in the classification when using all the features for classification. We then compared the classification accuracy using all the features and part of the features by conducting a paired t-test. It showed that there is minimal accuracy cost when including fewer features and there was no cures of dimensionality.

##### 4.6.7.2. Confusion matrix of classification analysis

The classification accuracy allowed us to check whether we can correctly classify the network labels of parcels. However, the classification accuracy alone can hide the detail we need to diagnose the performance of our model. For example, for multi-class classification, high classification accuracy may be observed because all classes are being predicted equally well or because one or two classes are being neglected by the model. Therefore, to better understand the performance of the classification model, we analyzed the confusion matrix, which is a summary of prediction results. In the confusion matrix, a row represents an instance of the actual class (i.e., an actual network), whereas a column represents an instance of the predicted class (i.e., the predicted network). The diagonal elements represent the number of points for which the predicted label is equal to the true label, while off-diagonal elements are those that are mislabeled by the classifier. Higher diagonal values indicate a higher number of correct predictions. The confusion matrix shows the ways in which our classification model is confused when it makes predictions. It gives us insight not only into the errors being made by the classifier but more importantly the types of errors that are being made. Specifically, we could explore the network similarity by analyzing the classification output and how the network similarity varies with task.

Due to the imbalance of observations across the networks, we normalized the confusion matrices by the number of elements in each class. In the current study, we only reported the normalized confusion matrix of control networks. We examined whether the percentages of predicted networks were different by conducting paired t-tests. We conducted FDR correction at p = 0.05 to control for multiple comparisons.

#### 4.6.8. Constructing feature similarity matrices

To investigate how FPCN subnetworks interact with DAN and DMN, we calculated the task and resting-state feature similarity. For each normalized feature matrix, we calculated Pearson correlation coefficients and transformed them to fisher z to represent the pairwise feature similarity between the time-series features of all possible combinations of brain parcels. As a result, a 400 × 400 feature similarity matrix was constructed for each individual and each run, representing the strength of the similarity of the local temporal fingerprints of brain areas. Finally, we averaged the estimates of feature similarity within networks, and between pairs of networks, to construct a network-by-network feature similarity matrix. The same method was used to calculate the resting-state feature similarity of the HCP dataset and construct a network-by-network feature similarity matrix.

#### 4.6.9. Constructing fMRI redundancy matrices

To investigate how FPCN subnetworks interact with DAN and DMN, we further calculated the redundant interaction. We used the recently developed information approach, integrated information decomposition, to decompose time-delayed mutual information into redundant, unique, and synergistic information shared with respect to both past and present state of both regions^37,43^. The redundant interaction quantifies how much information about the brain’s future trajectory is carried redundantly by distinct brain regions. We focused on the temporally persistent redundancy, which corresponds to redundant information in the past of both parts that is present in the future of both parts. The temporally persistent redundancy was calculated using the Gaussian solver implemented in the JIDT toolbox, based on their HRF-deconvolved BOLD signal time series. We calculated the redundant interaction for each pair of brain regions, resulting in a 400 × 400 redundancy matrix for per participant, per task and per run. Finally, we averaged the estimates of redundancy within networks, and between pairs of networks, to construct a network-by-network redundancy matrix, akin to the one that we constructed from the feature similarity data.

#### 4.6.10. Constructing fMRI functional connectivity matrices

To investigate how FPCN subnetworks interact with DAN and DMN, we further calculated the task and resting-state functional connectivity. We did not use the traditional psychophysiological interaction (PPI) to measure task-state functional connectivity because this method can inflate activation-induced task-state functional connectivity (i.e., identify regions that are active rather than interacting during the task) ^126^. Since task activations produce spurious but systematic inflation of task-based functional connectivity estimates ^126^, it is necessary to correct for the task-timing confounds by removing the first-order effect of task evoked activations (i.e., mean evoked task-related activity; likely active during the task) prior to estimating task-state functional connectivity (likely interacting during the task). Specifically, we fitted the task timing for each task using a finite impulse response (FIR) model ^126^. This method has been widely used before ^127–129^. We used an FIR model instead of a canonical hemodynamic response function or psychophysiological interactions (PPIs) given recent evidence suggesting that the FIR model reduces both false positives and false negatives when estimating functional connectivity ^126^.

After task regression, we demeaned the residual time series for each parcel and calculated the Pearson correlation as the functional connectivity for per participant, per task and per run. The Pearson correlation coefficients might be inflated due to the temporal autocorrelation present in task fMRI time series data ^130^. To account for the potential inflation of the Pearson correlation coefficients, we corrected the Pearson correlation using a novel correction approach – xDF. xDF accounts for distinct autocorrelation in each time series for instantaneous and lagged cross-correlation ^131^. We calculated xDF-adjusted z-scored correlation coefficients to compute interregional relationships of bold time series, resulting in a 400 × 400 functional connectivity matrix for per participant, per task and per run. Finally, we averaged the estimates of functional connectivity within networks, and between pairs of networks, to construct a network-by-network functional connectivity matrix. The same method was used to calculate the resting-state functional connectivity of the HCP dataset and construct a network-by-network functional connectivity matrix except that the task regression was not performed.

#### 4.6.11. Comparing feature similarity, redundancy and functional connectivity difference between networks

To investigate whether FPCN-A showed greater feature similarity with DAN than FPCN-B did in each task, we calculated the average feature similarity between FPCN-A and DAN and the feature similarity between FPCN-B and DAN, respectively across all runs per participant per task, then calculated the relative feature similarity difference (i.e., the feature similarity between FPCN-A and DAN minus the feature similarity between FPCN-B and DAN), and finally conducted paired-t tests for each task. Then we examined the feature similarity between FPCN-B and DMN versus FPCN-A and DMN in the same way. We conducted FDR correction at p = 0.05 to control for multiple comparisons. Similarly, we investigate whether FPCN-A showed stronger redundancy and functional connectivity with DAN than FPCN-B did. We also examined whether FPCN-B showed greater redundancy and functional connectivity with DMN than FPCN-A did. The procedure as above, except we extracted the redundancy and functional connectivity matrix, respectively.

We further investigated whether the task influenced the feature similarity difference between network pairs using the maximum/minimum permutation test. We might expect that feature similarity between FPCN-A and DAN-A versus FPCN-A and FPCN-B would be greater in spatial working memory task than in the association task because FPCN-A might shift more to DAN in the spatial working memory task that requires an external goal but no memory retrieval. To examine this possibility, we calculated the mean feature similarity difference between FPCN-A and DAN-A versus FPCN-A and FPCN-B for each task and calculated the mean feature similarity difference between each task pair. To test for statistical significance, we permutated the task label 10000 times; we then calculated the mean feature similarity difference between these two tasks to build a null distribution for each task pair. Since we included multiple task pairs, we used the permutation-based maximum mean feature similarity difference and minimum mean feature similarity difference values in the null distribution for each task pair to control the family-wise error (FWE) rate (p = 0.05, FWE-corrected). To evaluate significance, if the observed mean difference value was positive, we counted the percentage of times that mean difference values in the maximum null distribution were greater than the observed ‘true’ mean difference values; by contrast, if the observed mean difference value was negative, we counted the percentage of times of mean difference values in the minimum null distribution were less than the observed ‘true’ mean difference values.

Similarly, we investigated whether the task influenced the redundancy and functional connectivity difference between network pairs. The procedure was as above, except we extracted the redundancy and functional connectivity from the network-by-network redundancy and functional connectivity matrix, respectively. We conducted FDR correction at p = 0.05 to control for multiple comparisons.

#### 4.6.12. Task fMRI univariate analysis

To reveal the functional differentiation between FPCN-A and FPCN-B, we examined whether they showed similar or opposite activation patterns. We identified regions that were activated or deactivated in the tasks by building a general linear model (GLM). We also examined regions where the neural responses were modulated by task difficulty. For semantic tasks, we included one task mean regressor and one demeaned parametric regressor of semantic rating. We examined how the neural responses were negatively modulated by feature similarity rating for the matching trials and positively modulated by feature similarity rating for the non-matching trials. For the semantic association task, we examined how the neural responses were negatively modulated by semantic association strength rating for the related trials and positively modulated semantic association strength rating for the non-related trials. For the non-semantic tasks, we included two regressors – easy and hard conditions to reveal the regions showing greater activation in the hard than easy conditions. For all the tasks, we also modelled incorrect trials as regressors of no interest.

We extracted the beta value of each parcel in these task conditions and tested whether they were significantly activated (i.e., above zero) or deactivated (i.e., below zero) relative to implicit baseline (i.e., fixation period) to explore the task mean effect. Then we considered parcels that showed stronger or weaker activation in the hard than in the easy condition in the spatial working memory and math tasks and that the activations were positively modulated to semantic difficulty. These parcels were thought to support general executive control. Fixed-effects analyses were conducted using nilearn ^100^ to estimate the average effects across runs within each subject for each parcel. Then we conducted one-sample t-tests to assess whether the estimated effect-size (i.e., contrast) was significantly different from zero across all subjects. We conducted FDR correction at p = 0.05 to control for multiple comparisons. Finally, we identified the network that each parcel belonged to ^30^.

### 4.7. Data and materials availability

The HCP data is publicly available here https://www.humanconnectome.org/. The York data is not available due to insufficient consent. Researchers wishing to access the data should contact Elizabeth Jefferies or the Chair of the Research Ethics Committee of the York Neuroimaging Centre. Data will be released when this is possible under the terms of the UK GDPR. Analysis code for this study is available at https://github.com/Xiuyi-Wang/Control_Project.

## Acknowledgements

We are grateful to Pradeepa Ruwan and Antonia De Freitas for piloting the experiment. We are grateful to Andrea I. Luppi and Emmanuel A. Stamatakis for providing code to calculate redundancy. We would like to thank Ben D Fulcher for providing codes and guide to help us extract the features using hctsa. We would like to thank Ting Xu and colleagues for making available their data pertaining cortical areal expansion and cross-species similarity. We thank Nan Lin and Yanni Cui for helpful feedback on an earlier draft of our manuscript.

X. W. discloses support for the research of this work from Scientific Foundation of Institute of Psychology, Chinese Academy of Sciences (Grant Number. E1CX4725CX). Y.D. discloses support for the publication of this work from the STI 2030—Major Projects (Grant Number. 2021ZD0201500), the National Natural Science Foundation of China (Grant No. 31822024) and the Strategic Priority Research Program of Chinese Academy of Sciences (Grant Number. XDB32010300), and Scientific Foundation of Institute of Psychology, Chinese Academy of Sciences (Grant Number. E2CX3625CX). E. J. discloses support for the research of this work from a European Research Council Consolidator grant (Project ID: 771863 – FLEXSEM) to E.J.

