## Supplementary Material for "The brain’s topographical organization shapes dynamic interaction patterns to support flexible behavior"

Full title: The brain's topographical organization shapes dynamic coupling patterns to support flexible behavior

Short title: Cortical topography underpins flexible behavior

### 1.1. The homogeneity within MS-HBM-based individualized parcels was greater than that in the canonical Yeo 17-network group atlas.

We found that the homogeneity within multi-session hierarchical Bayesian model (MS-HBM) based individualized parcels was greater than that in the canonical Yeo 17-network group atlas that does not consider variation in functional neuroanatomy for each task (Fig. S1). The homogeneity of the semantic tasks in the York dataset (including semantic feature matching and semantic association tasks) was greater than that in the non-semantic tasks of York dataset (including spatial working memory and math tasks) (Figure S1). The homogeneity difference might be explained by several differences between the two datasets. Firstly, the data quality might be higher for the semantic than non-semantic tasks. T2w imaging was collected for semantic dataset but not for the non-semantic dataset. The high resolution T2w image can assist in pial surface placement. The blood vessels and dura are close to isointense with gray matter in the T1w image, however, they are clearly of different intensity in the T2w image, allowing them to be easily excluded from the gray matter ribbon. The inclusion of T2w image would reduce errors and improve quality control which was critical for performing the analysis on the surface ribbon. Secondly, the semantic tasks lasted longer than the non-semantic tasks and had more data. Each semantic task had 4 runs and each run lasted for about 10 mins, whereas each non-semantic task had 2 runs and the spatial working memory and math tasks lasted for 7min 28s and 5min 16s, respectively. The longer scanning time for the semantic tasks would improve the robustness of the functional connectivity pattern used for generating the individual-specific parcellation. Therefore, the precision of individual-specific parcellation is likely to be greater for the semantic than the non-semantic tasks.

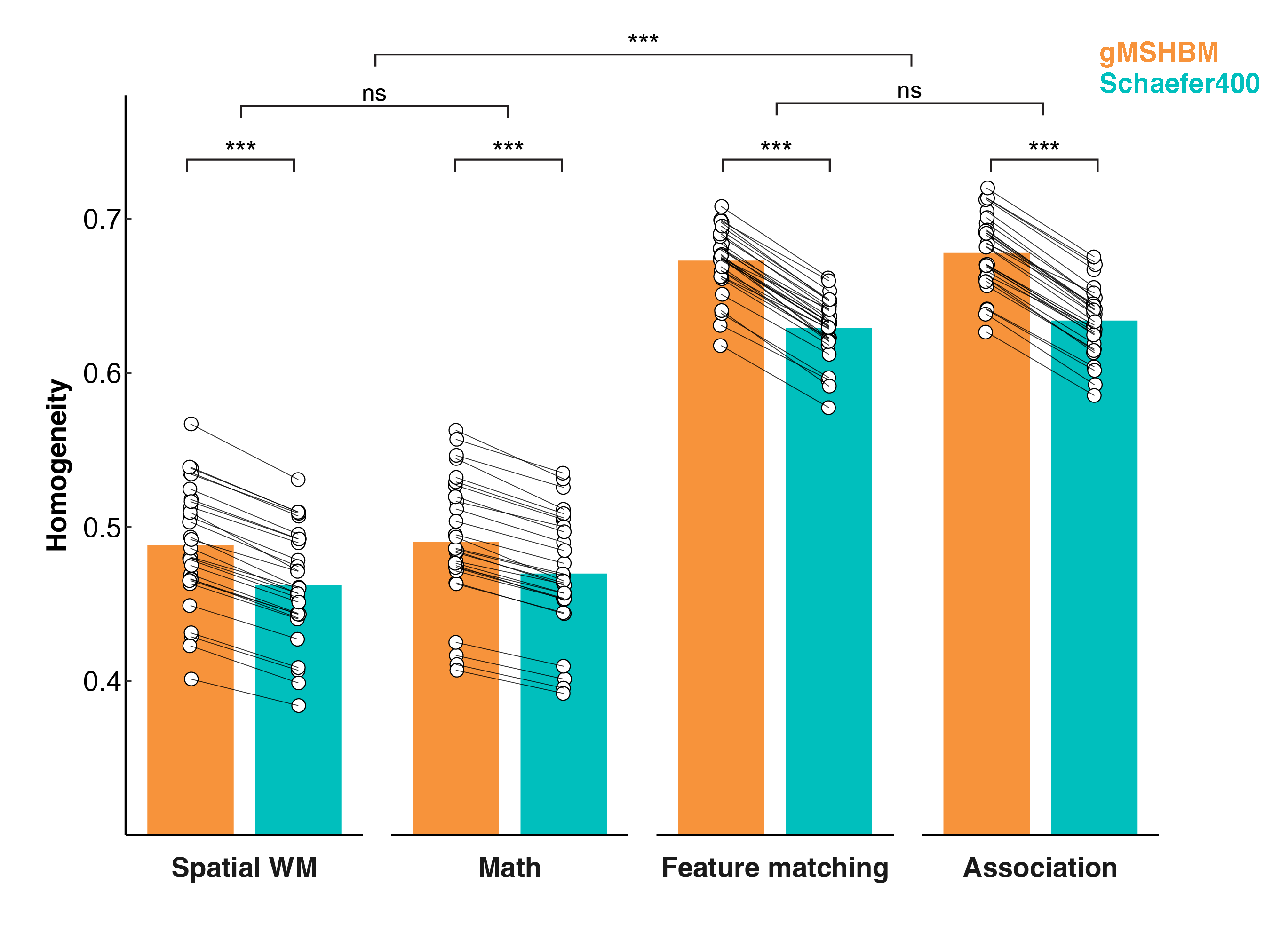

#### Fig S1. The homogeneity within MS-HBM-based individualized parcels was greater than that in the canonical Yeo 17-network group atlas that does not consider variation in functional neuroanatomy for each task. The homogeneity of the semantic tasks from the York dataset (including semantic feature matching and semantic association tasks) was greater than the non-semantic tasks (including spatial working memory and math tasks).

### 1.2. The network labels could be correctly predicted using the extracted features.

Classification accuracy was significantly greater than the chance level per participant per task, demonstrating that the classifier could correctly classify the network labels (rest: mean = 0.37, sd = 0.04; spatial working memory: mean = 0.24, sd = 0.03, math: mean = 0.21, sd = 0.04; semantic feature matching: mean = 0.38, sd = 0.03; semantic association: mean = 0.37, sd = 0.03, Fig. S2). The classification accuracy for rest and the semantic tasks was greater than that for the non-semantic tasks (p < 0.05, FWE corrected). This might be explained by the factors potentially contributing to the homogeneity difference between semantic and non-semantic tasks discussed above: higher data quality for the semantic tasks would ensure better consistency in the extracted features; higher homogeneity for these tasks suggests data is less likely to be mixed across parcels and networks; and more runs would provide more data for training and testing that would improve the classification accuracy. The classification accuracy difference between semantic tasks and non-semantic tasks means it is not meaningful to compare the confusion matrix across tasks. Below, we show the full confusion matrix for each task. The values on the diagonal (i.e., the percentage of predictions that were correct) were greater than the values off the diagonal (i.e., the percentage of wrong predictions), confirming that network labels could be correctly predicted. Generally, sensory-motor cortex had higher classification accuracy than transmodal cortex.

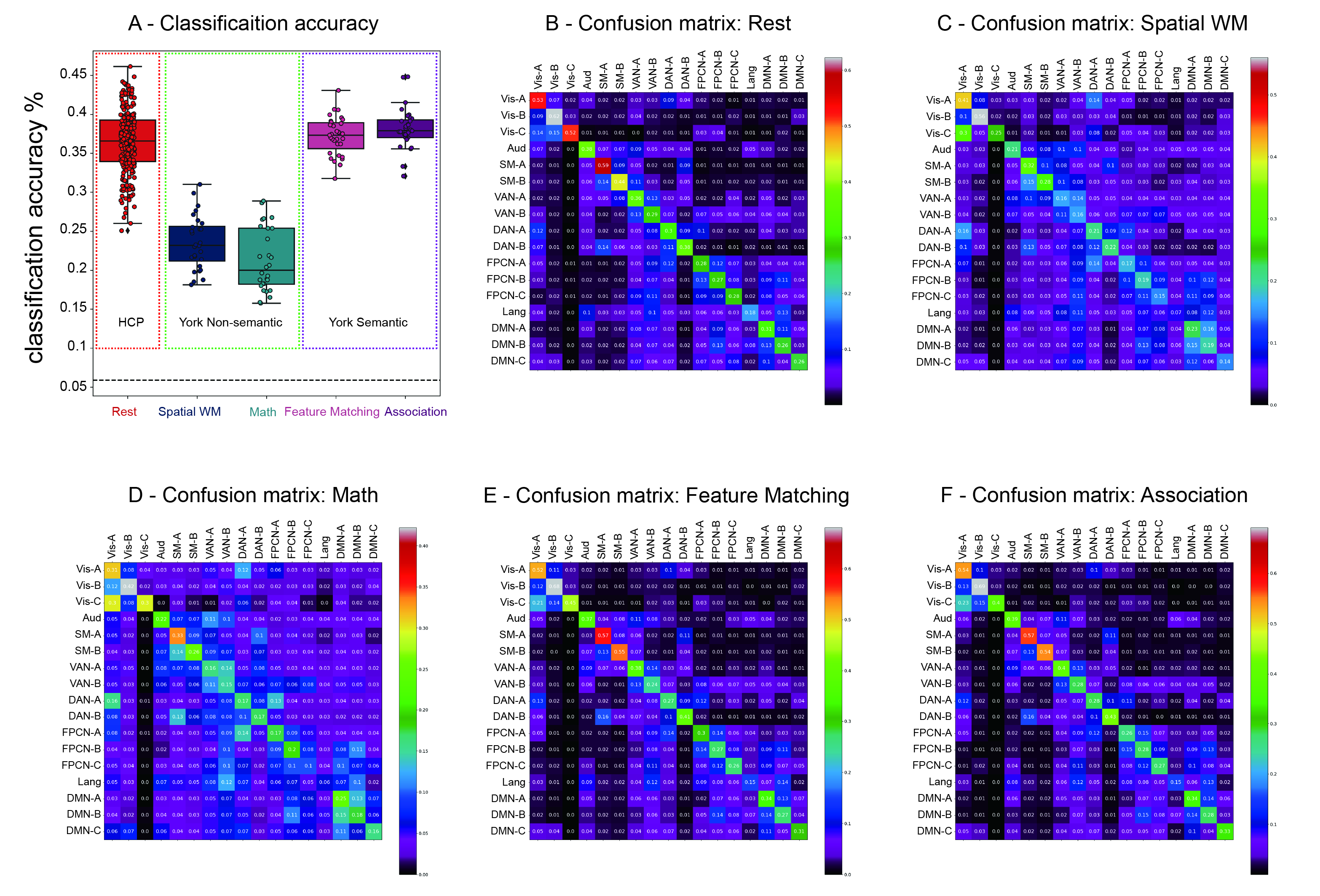

#### Fig S2. The network labels of parcels could be correctly predicted using the extracted feature vectors. A - Classification accuracy was significantly greater than the chance level per participant per task. The classification accuracy of the rest and semantic tasks was greater than that of the non-semantic tasks. The dashed line represents the chance level. B, C, D, E, F – The confusion matrix of the classification at rest, and in the spatial working memory, math, semantic feature matching, semantic association tasks, respectively.

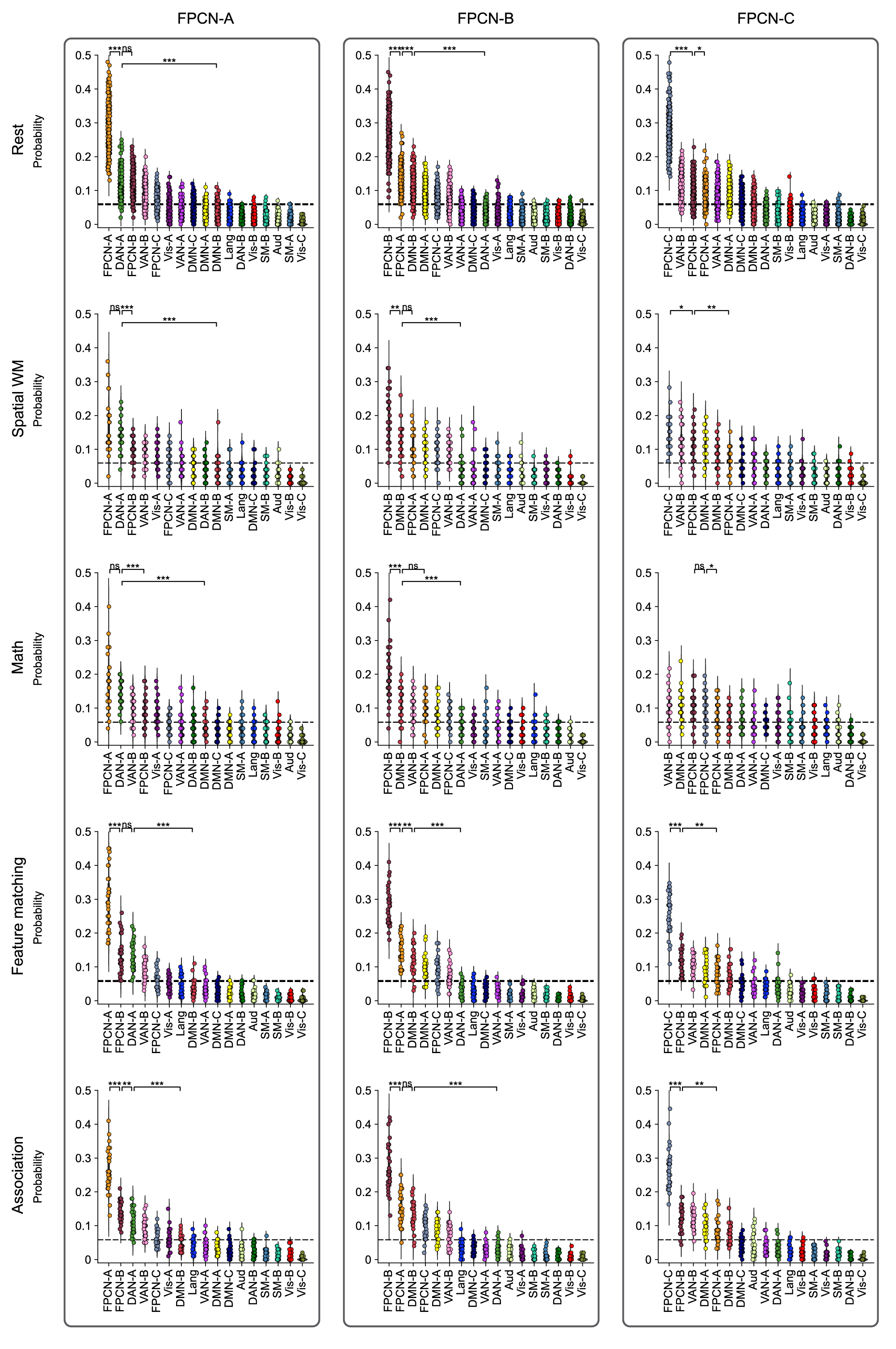

#### Fig S3. The probabilities of classifying FPCN-A, FPCN-B, and FPCN-C as themselves or misclassifying them as other networks. Left panel: FPCN-A was most likely to be incorrectly classified as DAN-A at rest and in non-semantic tasks, and as FPCN-B in semantic tasks. It was also misclassified as VAN-B, and not often misclassified as DMN-B. The probability of misclassifying FPCN-A as DAN-A was greater than as DMN-B and as VAN-B at rest and in all the tasks. Middle panel: FPCN-B was most likely to be misclassified as DMN-B in non-semantic tasks and as FPCN-A in semantic tasks and at rest. It was also misclassified as DMN-A, and not often misclassified as DAN-A. The probability of misclassifying FPCN-B as DMN-B was greater than as DAN-A and as VAN-B at rest and in all the tasks. Right panel: FPCN-C was most likely to be misclassified as VAN-A at rest, in the spatial working memory tasks, as FPCN-B in semantic tasks, and as DMN-A in the math tasks. The probability of misclassifying FPCN-C as VAN-B was greater than as DAN-A and DMN-B at rest and in all the tasks. The dashed line represents the chance level.

#### For completeness, we also ordered the networks by redundancy and functional connectivity with each FPCN subnetwork (Fig. S4 and Fig. S5).

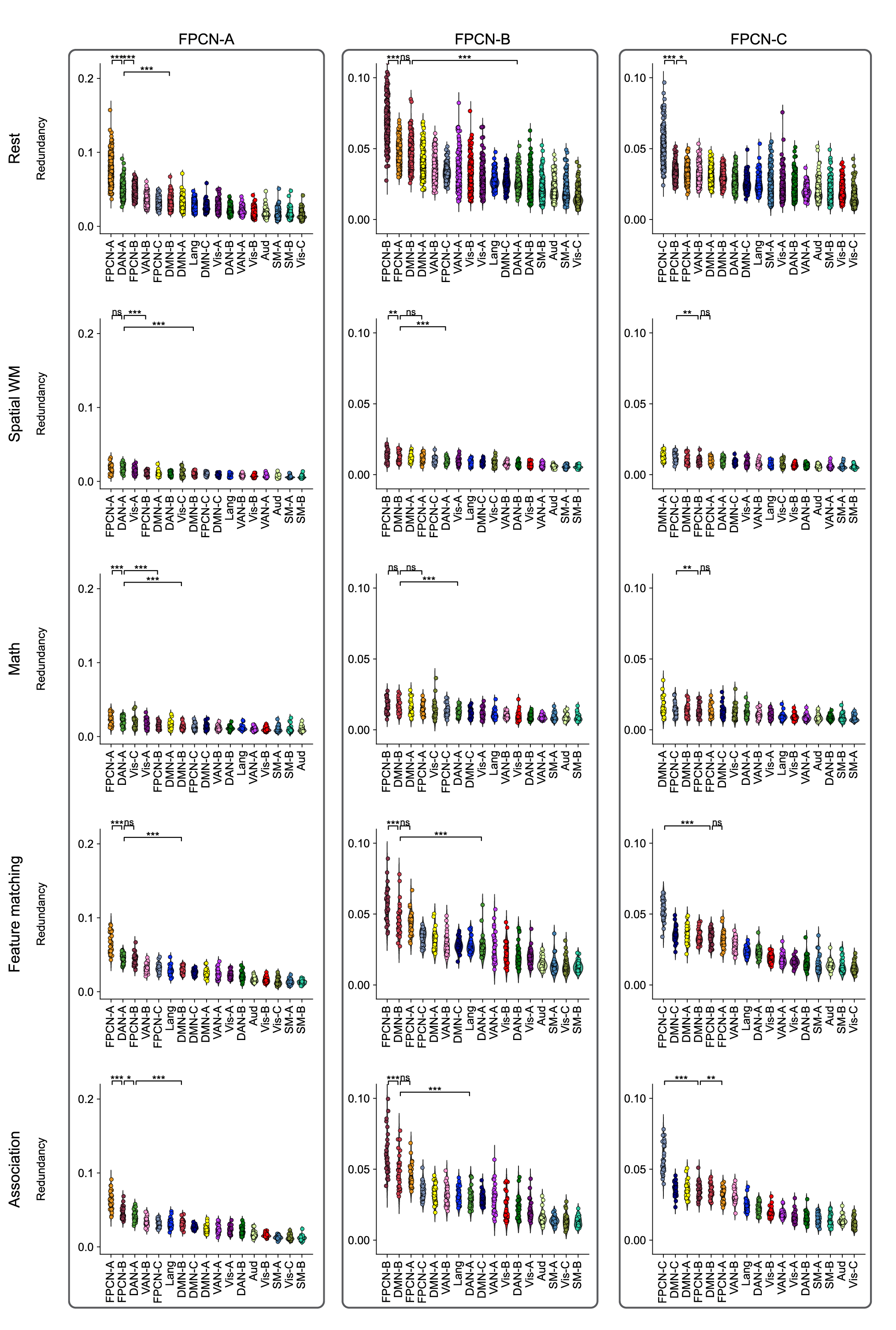

#### Fig S4. The redundant interaction (i.e., shared information) of FPCN subnetworks. The left panel shows the redundant interaction of FPCN-A. The top three networks showing highest redundant interaction were often FPCN-A, FPCN-B and DAN-A. FPCN-A showed greater redundant interaction with the visual network in the non-semantic tasks, but this effect was not observed in the semantic tasks although all the stimuli were visually presented. FPCN-A always showed greater redundancy with DAN-A than with DMN-B. Middle panel shows the redundant interaction of FPCN-B. The top three networks showing highest redundant interaction were often FPCN-B, FPCN-A and DMN-B. FPCN-B always showed greater redundancy with DMN-B than with DAN-A. The right panel shows the redundant interaction of FPCN-C. Unlike FPCN-A and FPCN-B which showed clear redundant interaction patterns at rest and in all the tasks, the pattern of FPCN-C was complex.

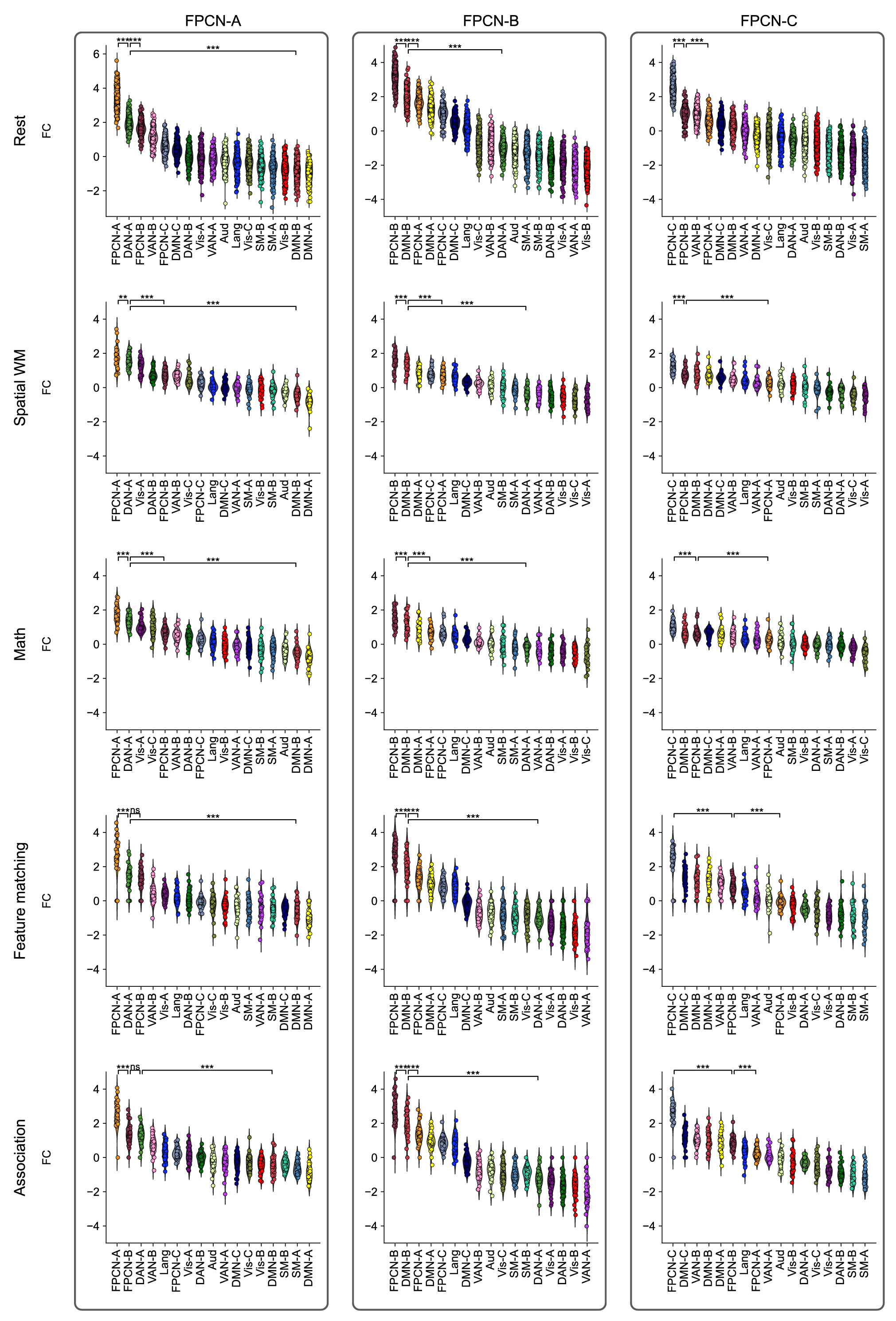

#### Fig S5. The functional connectivity (FC) patterns of FPCN subnetworks. The left panel shows the functional connectivity of FPCN-A. The top three networks showing highest functional connectivity with this network were often FPCN-A, FPCN-B and DAN-A. FPCN-A always showed greater functional connectivity with DAN-A than with DMN-B. Middle panel shows the functional connectivity of FPCN-B. The top three networks showing highest functional connectivity with this network were often FPCN-B, FPCN-A and DMN-B. FPCN-B always showed greater functional connectivity with DMN-B than with DAN-A. The right panel shows the functional connectivity of FPCN-C.

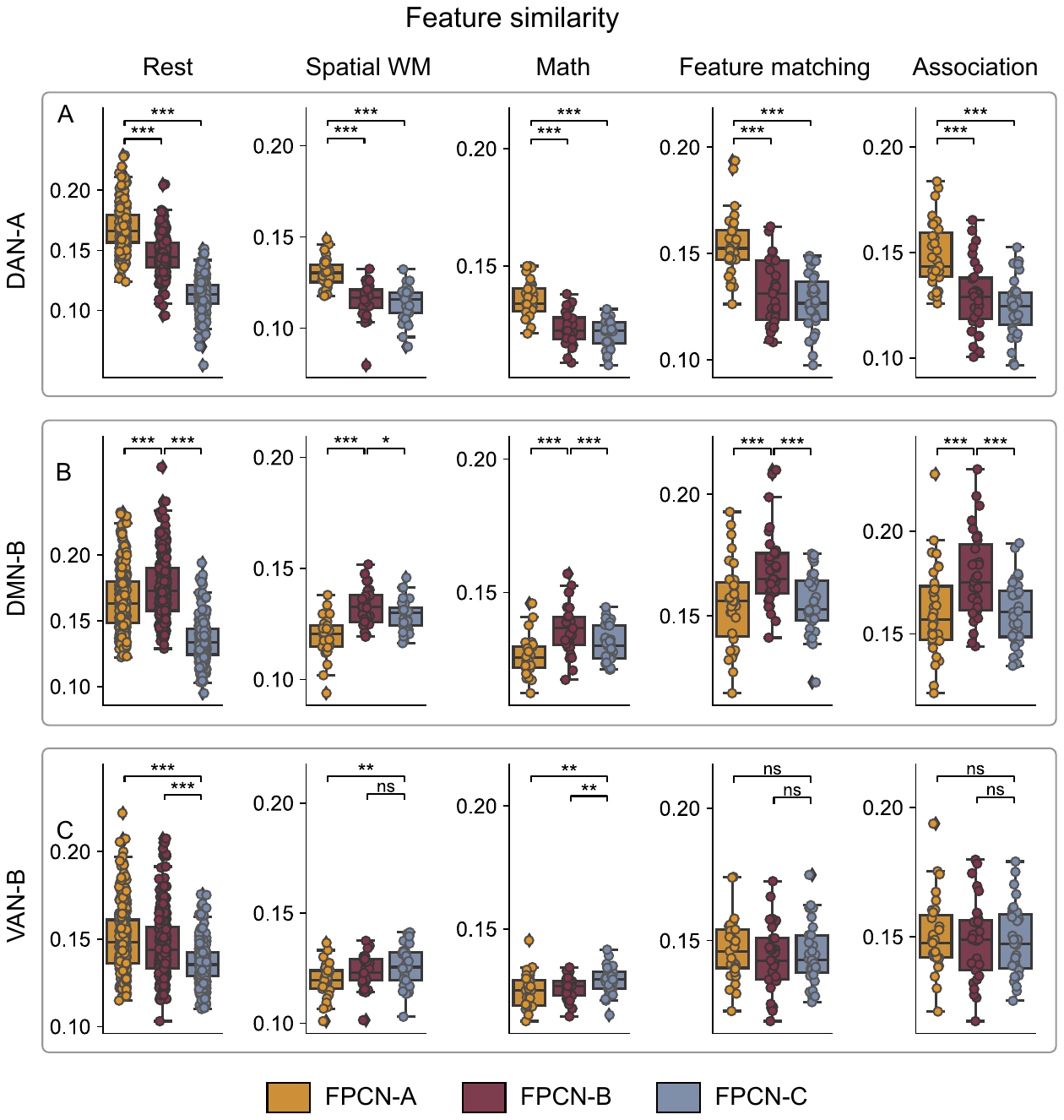
Fig S6. FPCN-A, FPCN-B and FPCN-C coupled with different networks. FPCN-A always coupled with DAN-A while FPCN-B always coupled with DMN-B across different conditions. But FPCN-C does not always couple with a specific network. Upper panel: FPCN-A showed stronger feature similarity with DAN-A than FPCN-B and FPCN-C did. Middle panel: FPCN-B showed stronger feature similarity with DMN-B than FPCN-A and FPCN-C did. Bottom panel: FPCN-C does not always show stronger feature similarity with VAN-B than FPCN-A and FPCN-B. FPCN = fronto-parietal control network. DAN = dorsal attention network, DMN = default mode network, WM = working memory.

### 1.3. FPCN-A and DAN-A showed similar activation patterns, while FPCN-B and DMN-B showed similar deactivation patterns.

After showing greater coupling of FPCN-A with DAN-A and FPCN-B with DMN-B, we examined whether the same network pairs showed similar activation patterns across tasks. We examined task activation and deactivation. Most parcels in FPCN-A and DAN-A were active across all the tasks although several parcels showed deactivation in the semantic tasks (Fig. S7). FPCN-B and DMN-B showed a different pattern, with more deactivation across the tasks. Only three parcels of FPCN-B showed task activation (Fig. S7).

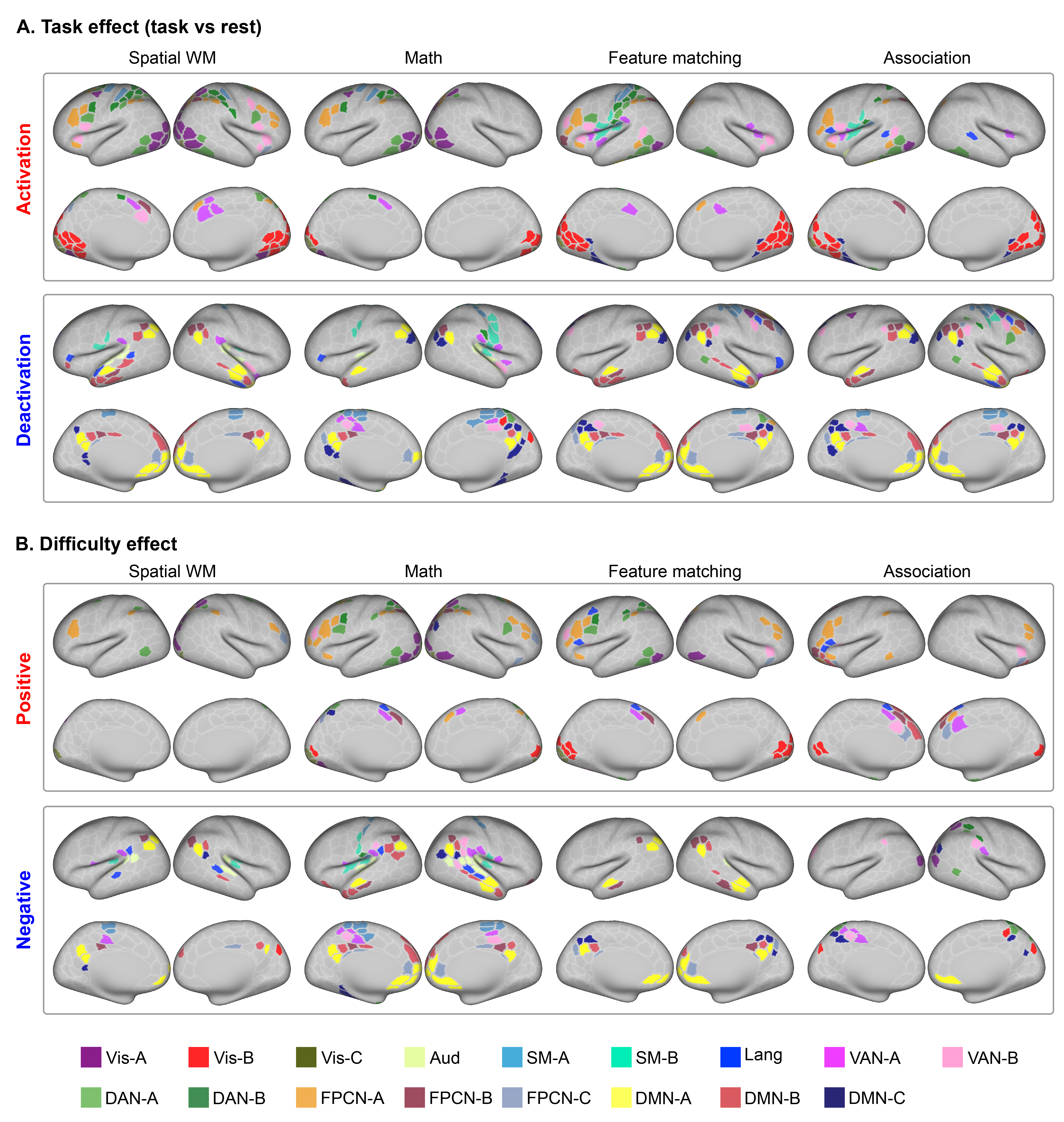

Fig S7: FPCN-A and DAN-A showed task activation, while FPCN-B and DMN-B showed task deactivation. A: Regions showing activation and deactivation relative to the implicit baseline (i.e., rest) when doing the tasks. B: Top panel - Regions showing stronger activation in hard than easy conditions in the non-semantic tasks, including spatial working memory and math tasks. Regions whose response were positively modulated by the feature similarity rating in the semantic feature matching and semantic association strength rating in the semantic association tasks. Bottom panel - Regions showing stronger activation in easy than hard conditions in the non-semantic tasks, including spatial working memory and math tasks. Regions whose response were negatively modulated by the feature similarity rating in the semantic feature matching and semantic association strength rating in the semantic association tasks.

1.4. FPCN-A shared more information with DAN-A than with FPCN-B in the non-semantic tasks but showed the opposite pattern in the semantic tasks.

Next, we compared the difference in redundancy between FPCN-A and DAN-A versus FPCN-A and FPCN-B within each task and rest. As shown in Fig. 6A, FPCN-A showed greater redundant interaction with DAN-A than with FPCN-B at rest (t = 3.645, p < 0.001) and in the non-semantic tasks (spatial working memory task, t = 8.141, p < 0.001; math task, t = 6.874, p < 0.001) but equal redundant interaction in the semantic feature matching task (t = 0.316, p = 0.754). FPCN-A showed less redundancy with DAN-A than with FPCN-B in the semantic association task (t = -2.190, p = 0.045). All p values are FDR-corrected.

### 1.5. FPCN-A showed greater functional connectivity with DAN-A than with FPCN-B in the non-semantic tasks but showed the opposite pattern in the semantic tasks.

We compared the difference in functional connectivity between FPCN-A and DAN-A versus FPCN-A and FPCN-B within each task and rest. As shown in Fig. 6B, FPCN-A showed stronger functional connectivity with DAN-A than with FPCN-B at rest (t = 5.892, p < 0.001). The same pattern was seen in the non-semantic tasks (spatial working memory task, t = 11.478, p < 0.001; math task, t = 7.146, p < 0.001), but no differences were observed in the semantic tasks (feature matching task, t = 0.698, p = 0.490; association task, t = 0.759, p = 0.454). All p values are FDR-corrected.

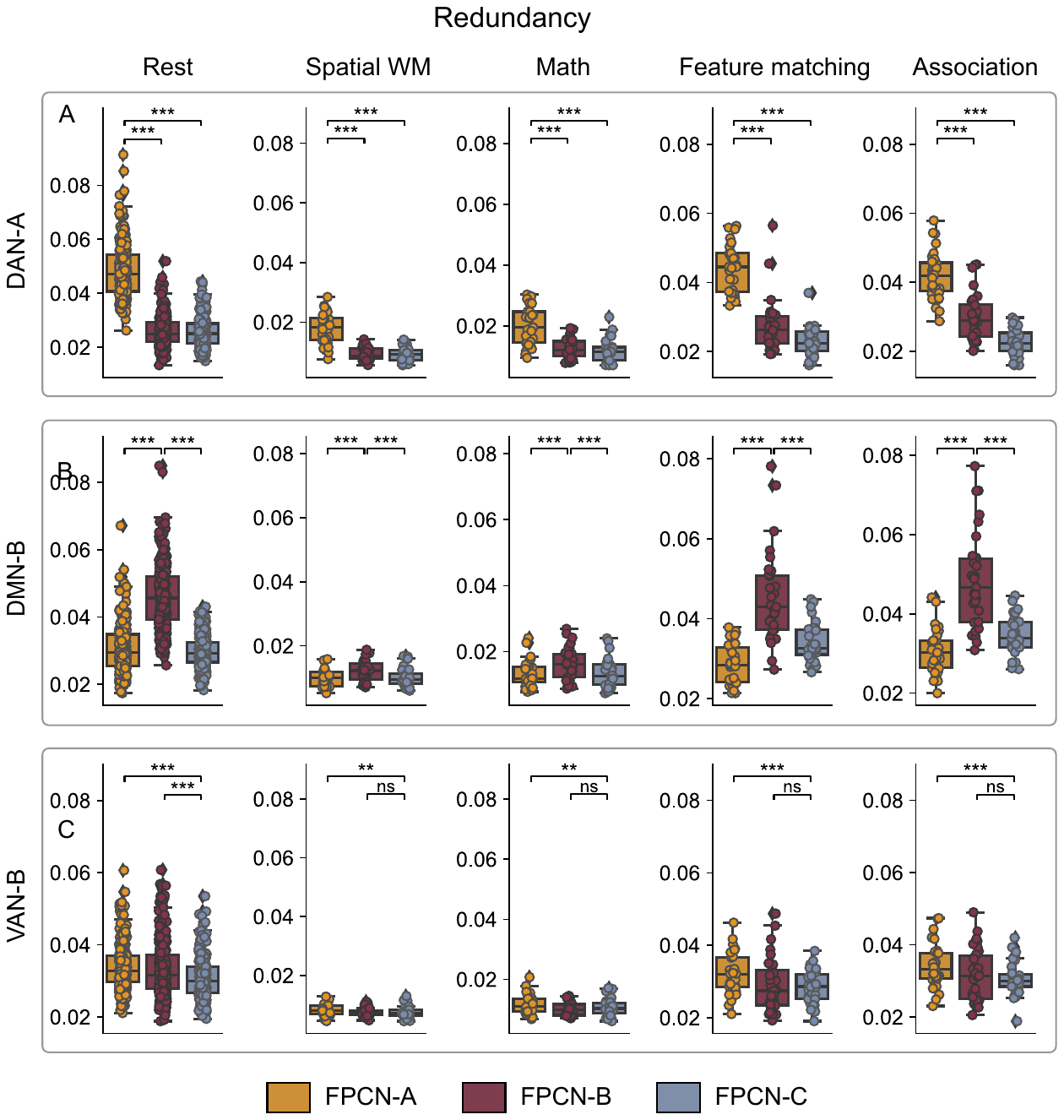

#### Fig S8: Top panel - Compared with FPCN-B and FPCN-C, FPCN-A showed greater redundancy with DAN-A. Middle panel - Compared with FPCN-A and FPCN-C, FPCN-B showed greater redundancy with DMN B. Bottom panel – FPCN-C did not always show greater redundancy with VAN-B.

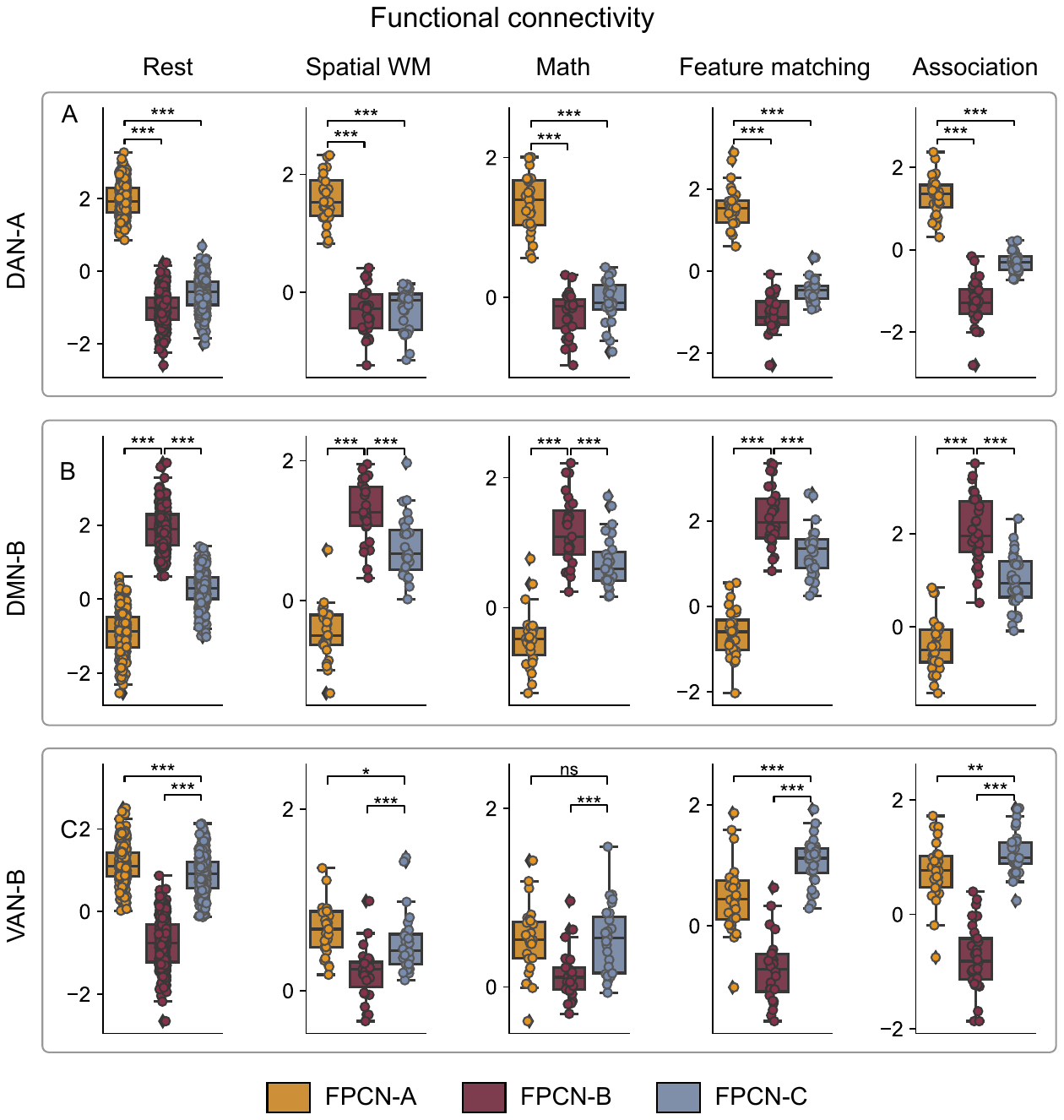

#### Fig S9: Top panel - Compared with FPCN-B and FPCN-C, FPCN-A showed stronger functional connectivity with DAN-A. Middle panel - Compared with FPCN-A and FPCN-C, FPCN-B showed stronger functional connectivity with DMN B. Bottom panel – FPCN-C did not always show greater functional connectivity with VAN-B.

#### Table S1 FPCN-A was more likely to be misclassified as DAN-A than as DMN-B, while FPCN-B showed the opposite pattern.

| True network | Task | Mean probability | | | DAN-A vs DMN-B | | |
| --- | --- | --- | --- | --- | --- | --- | --- |
|  |  | DAN-A | DMN-B | *t* | | *p* | *q* |
| FPCN-A | Rest | 0.118 | 0.040 | 27.313 | | 3.66E-76 | 1.83E-75 |
|  | Spatial WM | 0.144 | 0.043 | 9.191 | | 8.41E-10 | 1.05E-09 |
|  | Math | 0.143 | 0.044 | 9.887 | | 1.81E-10 | 3.02E-10 |
|  | Feature matching | 0.132 | 0.036 | 10.908 | | 5.82E-12 | 1.45E-11 |
|  | Association | 0.111 | 0.047 | 7.087 | | 7.01E-08 | 7.01E-08 |
| FPCN-B | Rest | 0.036 | 0.111 | -26.184 | | 8.00E-73 | 4.00E-72 |
|  | Spatial WM | 0.042 | 0.116 | -5.070 | | 2.52E-05 | 2.52E-05 |
|  | Math | 0.044 | 0.111 | -5.516 | | 7.64E-06 | 9.55E-06 |
|  | Feature matching | 0.032 | 0.115 | -10.358 | | 2.00E-11 | 3.34E-11 |
|  | Association | 0.027 | 0.128 | -14.129 | | 8.50E-15 | 2.12E-14 |

q is the FDR-corrected p value.

#### Table S2 – DAN-A showed greater redundancy with FPCN-A than with FPCN-B; DMN-B showed the opposite pattern.

| Network | Task | Mean Redundancy | | FPCN-A vs FPCN-B | | |
| --- | --- | --- | --- | --- | --- | --- |
|  |  | FPCN-A | FPCN-B | *t* | *p* | *q* |
| DAN-A | Rest | 0.048 | 0.026 | 38.033 | 2.32E-75 | 1.53E-104 |
|  | Spatial WM | 0.018 | 0.009 | 9.360 | 2.51E-07 | 4.06E-10 |
|  | Math | 0.020 | 0.012 | 10.087 | 9.52E-10 | 7.95E-11 |
|  | Feature matching | 0.044 | 0.027 | 11.408 | 3.64E-12 | 1.95E-12 |
|  | Association | 0.042 | 0.030 | 9.544 | 2.67E-11 | 1.34E-10 |
| DMN-B | Rest | 0.031 | 0.046 | -21.934 | 1.57E-20 | 1.22E-59 |
|  | Spatial WM | 0.010 | 0.012 | -5.010 | 1.85E-07 | 2.70E-05 |
|  | Math | 0.013 | 0.016 | -4.714 | 1.32E-07 | 6.06E-05 |
|  | Feature matching | 0.029 | 0.045 | -9.071 | 8.05E-05 | 4.22E-10 |
|  | Association | 0.031 | 0.048 | -9.519 | 6.75E-07 | 1.42E-10 |

#### Table S3 – DAN-A showed greater functional connectivity (FC) with FPCN-A than with FPCN-B; DMN-B showed the opposite pattern.

| Network | Task | Mean FC | | FPCN-A vs FPCN-B | | |
| --- | --- | --- | --- | --- | --- | --- |
|  |  | FPCN-A | FPCN-B | *t* | *p* | *q* |
| DAN-A | Rest | 1.955 | -1.026 | 73.471 | 2.01E-167 | 2.01E-168 |
|  | Spatial WM | 1.589 | -0.326 | 17.545 | 8.79E-16 | 6.24E-16 |
|  | Math | 1.341 | -0.226 | 15.901 | 6.53E-15 | 6.53E-15 |
|  | Feature matching | 1.545 | -1.045 | 22.901 | 6.40E-19 | 3.20E-19 |
|  | Association | 1.299 | -1.262 | 22.413 | 1.99E-19 | 7.33E-20 |
| DMN-B | Rest | -0.893 | 1.893 | -64.832 | 4.12E-155 | 8.23E-156 |
|  | Spatial WM | -0.443 | 1.273 | -17.458 | 8.79E-16 | 7.03E-16 |
|  | Math | -0.484 | 1.175 | -17.281 | 9.98E-16 | 8.98E-16 |
|  | Feature matching | -0.610 | 2.100 | -24.169 | 1.99E-19 | 7.98E-20 |
|  | Association | -0.419 | 2.059 | -17.887 | 5.51E-17 | 3.31E-17 |
